# Hematopoietic stem cells are a reservoir for trained immunity in autoimmune disease

**DOI:** 10.1101/2022.05.20.492533

**Authors:** Taylor S. Mills, Bailee Kain, Matt A. Burchill, Etienne Danis, Rachel Culp-Hill, Andrew Goodspeed, Sarah Ferrara, James R. Roede, Angelo D’Alessandro, Katherine Y. King, Eric M. Pietras

## Abstract

Trained immunity (TI) has been speculated to serve as a contributor to autoimmune disease (AD) pathogenesis via generation of hyper-inflammatory myeloid cells. Using a mouse model of systemic lupus erythematosus (SLE), we show that hematopoietic stem cells (HSC) constitute a transplantable, long-term reservoir for macrophages that exhibit features of TI, including increased *Mycobacterium avium* killing, inflammatory cytokine production, and augmented capacity to co-stimulate naive T cells. Strikingly, hematopoietic progenitor cells derived from these HSC exhibit unique molecular features characterized by reduced chromatin accessibility and transcription of metabolic genes, accompanied by reduced glycolysis and central carbon metabolism. Altogether, our data identify HSC as a functional unit of TI in chronic AD, establish increased T-cell costimulatory activity as a potentially pathogenic feature of TI in this setting, and show that macrophages inherit reduced metabolic activity from AD-exposed HSC, suggesting metabolic activation and TI can be decoupled. Our findings thus implicate HSC as a long-term reservoir for TI that could contribute to AD.

## INTRODUCTION

Autoimmune diseases (AD) impact approximately 6% of the population (Davidson and Diamond, 2001; Wang, Wang and Gershwin, 2015; Theofilopoulos, Kono and Baccala, 2017) and rates are increasing globally (Lerner, Jeremias and Matthias, 2015; Dinse *et al*., 2020). While the complex etiology of these diseases are not completely understood, a hyperinflammatory environment generated by activated myeloid cells can lead to bystander activation of autoimmune lymphocytes (Laria *et al*., 2016; Navegantes *et al*., 2017; Morell, 2017; Ushio *et al*., 2017; Ma *et al*., 2019). This occurs when myeloid cells express inflammatory cytokines, and co-stimulatory ligands that can activate autoimmune T cells without TCR stimulation (Sfriso *et al*., 2010; Hussein and Rahal, 2019) leading to the development of AD. Thus, understanding mechanisms that potentiate the inflammatory activity of myeloid cells is crucial to understanding AD pathogenesis.

Trained immunity (TI) is described as a heightened response of myeloid innate immune cells to a secondary infection or stimuli due to memory of prior activation (Netea, Quintin and van der Meer, 2011). Treatment of humans, mice, or myeloid cells directly with inflammatory factors such as β-glucan and the Bacillus Calmette-Guerin vaccine (BCG) can trigger an antigen non-specific response of amplified inflammatory functions upon restimulation (Kleinnijenhuis *et al*., 2012; Arts *et al*., 2015, 2018; Bekkering *et al*., 2016; Kaufmann *et al*., 2018; Cirovic *et al*., 2020; Moorlag, Khan, *et al*., 2020; Moorlag, Rodriguez-Rosales, *et al*., 2020). This phenotype is characterized by increased generation of the inflammatory cytokines TNF-α, IL-6, and IL-1β, as well as increased capacity to kill live pathogens both *in vitro* and *in vivo*. Further, increased energy metabolism, mainly glycolysis, required to establish TI (Cheng *et al*., 2014; Arts *et al*., 2016; Domínguez-Andrés *et al*., 2017; Mitroulis *et al*., 2018). This in turn activates the TCA cycle which generates metabolites necessary to induce epigenetic programming associated with TI (Arts *et al*., 2016; Domínguez-Andrés *et al*., 2017, 2019). Interestingly, the time gap between the initial ‘training’ and the secondary stimulus can last well beyond the lifetime of individual myeloid cells (Kristensen, Aaby and Jensen, 2000; Garly *et al*., 2003; Vaugelade *et al*., 2004; Kleinnijenhuis *et al*., 2012; Arts *et al*., 2015). Thus, it has been speculated that hematopoietic stem and progenitor cells (HSPC) serve as a reservoir for trained immunity (Kaufmann *et al*., 2018; Mitroulis *et al*., 2018; Cirovic *et al*., 2020; de Laval *et al*., 2020).

Based on the above observations, we investigated whether HSC^LT^ are a reservoir for hyper-inflammatory trained macrophages. To test this hypothesis, we employed the pristane-induced mouse model of SLE and found that pristane-treated mice generate macrophages with key hallmarks of TI, including inflammatory and antimicrobial activity in response to live *Mycobacterium avium* infection. Using transplantation, we identify HSC^LT^ as the reservoir for TI. Interestingly, we find that HSC^LT^ from pristane-treated donors pass on an epigenetic program to myeloid progenitors characterized by reduced accessibility and transcription of metabolic genes, particularly in the mTOR and Myc pathways. Notably, while macrophages derived from these cells exhibit reduced central carbon metabolism upon stimulation, they retain key features of TI. Taken together, our data identify a key role for HSC^LT^ in TI that may contribute to AD pathology.

## MATERIALS AND METHODS

### Mice

6-8-week-old female C57BL/6J (Strain #000664) mice were obtained from Jackson laboratories. Mice were bred and maintained ventilated cages under specific pathogen free conditions in a temperature and light controlled environment with chow and water *ad libitum* for all experiments. For Lupus disease initiation 500μL of pristane oil (2,6,10,14-tetramethylpentadecane; Sigma Aldrich P9622) was injected intraperitoneal (i.p.), and age-gender-matched control animals received 500μL of sterile phosphate buffered saline. All animal procedures were approved by the University of Colorado Denver Anschutz Medical Campus Institutional Animal Care and Use Committee (IACUC).

### Flow cytometry and HSC isolation

Bone marrow isolation and cell staining was performed on ice in staining medium (SM; Hanks Balanced Salt Solution [HBSS 1x, Corning 1-022-CM] supplemented with heat inactivated fetal bovine serum [FBS, VWR 97068-05] to a final concentration of 2%). Femurs and tibiae were flushed with 3mL of SM and followed by RBC lysis with ACK lysis buffer. Viable cells were counted on a ViCELL (Beckman Coulter) and 1x10^7^ cells were stained for hematopoietic stem and progenitor cell (HSPC) analysis, and 1x10^6^ cells were stained for mature cell analysis. Splenocyte and thymocyte surface staining was performed by crushing and filtering the tissue through a 0.70μm cell strainer followed by RBC lysis. Flow cytometry was performed on the BD FACSCelestia or BD LSRII. For HSC isolation femur, tibiae, hips, humeri, and spines were crushed with a mortar and pestle in SM followed by RBC lysis and then separated using histopaque 1119 gradient centrifugation (Sigma, 11191). BM progenitor cells were enriched through c-Kit (CD117) positivity with c-Kit magnetic microbeads (130-091-224, Miltenyi) twice on a AutoMACS Pro (Miltenyi) before being stained for HSPC markers. BM cells were then double sorted to purity on a BD FACSAria Fusion at 20psi using a 100μm nozzle. For HSPC analysis cells were stained for 30 min on ice with the following antibodies: Lineage, PE-Cy-5-conjugated anti-CD3, CD4, CD5, CD8, Gr-1, B220, and Ter119; plus Sca-1-BV421, CD41-BV510, Flk2-Biotin, FcγR-BV711, CD150-BV786, CD34-FITC, EPCR-PE, Mac-1-PE/Cy7, ESAM-APC,

CD48-AlexaFluor 700, and c-Kit-APC/Cy7 (Biolegend), 50μg/mL Rat IgG (Sigma) was included as block with all staining. Cells were then incubated on ice for 30min before being washed with SM and centrifuged at 1200 x g, then stained with streptavidin-BV605 for 30 min on ice before another wash cycle. Cells were then resuspended in SM containing propidium iodide (PI) to stain any dead cells. For CLP and myeloid progenitor analysis cells were surface stained with the HSPC Abs cocktail with the following adjustments: Lineage, includes CD11b-PE/Cy5; CD105-BV786, CD150-PE, IL-7R-PE/Cy7. Mature cell populations were stained with the following antibodies: Gr-1-Pacific Blue, Ly6C-BV605, B220-BV786, CD4-FITC, CD8-PE, CD11b-PE/Cy7, IgM-APC, CD3-AlexaFluor700, CD19-APC/Cy7 (Biolegend).

### HSC transplantation

For transplantation experiments, congenic Boy/J (CD45.1) mice were lethally irradiated using a (split dose 11gy 3h apart) and were then transplanted via the retro-orbital vein with 500 LT-HSC or 2000 HSC plus 5x10^5^ Sca-1 depleted cells in a volume of 100μl of HBSS + 2% FBS. Recipient mice were then placed in ventilated cages with antibiotic water containing neomycin and polymixin for four weeks. Donor chimerism was determined at four-week intervals post transplantation using peripheral blood obtained from the submandibular vein.

### Generation of bone marrow derived macrophages

Bone marrow derived macrophages were generated by flushing one femur with 3mL of SM. BM cells were then placed into culture in T-75 flasks with BMDM media (DMEM base, 10% FBS, 1x Anti Anti (Gibco, 15240-062), 1x MEM-NEAA (Gibco, 11140-050), 1x Na-Pyruvate (Gibco, 11360-070), 1x L-glutamine [Gibco, 25030-081) containing 50ng/mL of M-CSF (Peprotech). On day 5 of culture, adherent cells were washed three times with sterile 1x DPBS (Gibco, 14190-136) and isolated via treatment with 0.25% Trypsin (Gibco, 25200-056) for 3 minutes at 37^oC^ prior to mechanical scraping. Cells were counted via hemocytometer before being seeded overnight prior to functional assessment. The positive generation of BMDMs in culture was determined by surface expression of Gr1+CD11b+F4/80+ via flow cytometry.

### BMDM *M. avium ex vivo* killing assays

Purified BMDMs were seeded into 6 or 12 well plates at concentration of 500,000 -1,000,000 or 200,000 cells/well, respectively. The following day, cells were either left untreated or infected with an MOI of 5 *M. avium*/seeded BMDM. Cells were incubated with bacteria for four hours or 3 days (protocol modified from Kaufmann *et al*., 2018). At harvest, supernatant from each well was collected for cytokine profiling and stored at -80C. Cells were then washed with 1 x PBS, scraped, and resuspended into Eppendorf tubes. Following manual counting using a hematocytometer, cells were spun down in a table-top centrifuge for ten minutes and resuspended in sterile dH_2_O to lyse eukaryotic cells. Cell suspensions were then frozen at -80^C^ until pre-amplification of DNA for qPCR.

A standard curve to determine *M. avium* concentration was generated using our laboratory stock of *M. avium*. An aliquot of bacteria was spun down at 13,000 RPM in a table-top centrifuge, resuspended in sterile dH_2_O, and boiled at 95C for 10 minutes to generate a stock of 16s DNA. Following boiling, DNA was quantified using a Nanodrop Microvolume UV-Vis Spectrophotometer (ThermoFisher Scientific) and aliquoted into serial dilutions ranging from undiluted DNA – 10^-6^. Standard curve samples were pre-amplified using Pre-Amp Mastermix (Fluidigm, San Francisco, CA) according to manufacturer’s instructions. Diluted pre-amplification product was used to complete RT-PCR using iTaq Universal SYBR Green Supermix (BioRad) and 16s *M. avium* primers. Ct values generated by these samples were then used as a standard to quantify *M. avium* copy number in samples from BMDM killing assays. BMDM killing assay samples were processed and ran using the same protocol as standard curve generation.

### BMDM T cell Proliferation assay

BMDMs were harvested and stimulated with the following conditions: BMDM media alone, or Immune complexes of Ovalbumin (50 mg/ml – Sigma, St. Louis, MO) and anti-OVA antibody (Biolegend, TOSGAA1) for 24 hours at 37°C and 5% CO_2_. Following the indicated stimulation BMDMs were washed 2x with Phosphate Buffered Saline (PBS, Corning Life Sciences, Corning, NY) and resuspended in RPMI 1640 (Gibco, Waltham, MA) supplemented with 10% Fetal Bovine Serum (Atlanta Biologicals, Flowery Branch, GA). BMDMs were then enumerated using a TC20 automatic cell counter (BioRad, Hercules, CA). To measure the ability of BMDMs to present peptide to antigen specific T cells, 20,000 BMDMs were plated in a 96 well plate with 50,000 purified naive OT-I T cells labeled with Violet Proliferation Dye 450 (BD Biosciences, Franklin Lakes, NJ). Following 72 hours of incubation at 37°C and 5% CO_2_, cells were harvested, stained with aCD8a (53-6.7) and aCD44 (IM7) from Bolegend (San Diego, CA) and samples were acquired on a BD FACSCanto II (BD Biosciences). Analysis of flow cytometry data was performed using FlowJo software V10.8.0 (Tree Star, Ashland, OR). OTI percent divided and division index was calculated as previously described (Roederer, 2011).

### Cytokine Analysis

Cellular supernatant was isolated from the *ex vivo* cultures prior to cellular harvest and frozen at –80^C^ until further use. Samples were shipped overnight on dry ice to Eve technologies (Calgary, Canada) and assessed using the Mouse Cytokine Array Proinflammatory Focused 10-plex (MDF10) for cytokine release assessment of macrophages.

### Seahorse extracellular flux assays

Prior to plating, the cell culture plate was treated with Cell-Tak™ (Corning 35424) according to the manufacturers protocol. Then, 75,000 BMDMs were plated and allowed to adhere overnight using the BMDM media. For wells that were stimulated with immune complexes (ImmCom), BMDMs were treated with ImmCom at the time of plating. The following day the Seahorse extracellular flux cartridge (Agilent 102601-100) was prepared according to manufacturer’s instructions with the XF calibrant. For the mitochondrial stress test, we used Oligomycin (20μM, 20μL port A; Sigma 75351), FCCP (20μM, 22μL port B; Sigma C2920), and Rotenone/Antimycin-A (5μM, 25μL port C; Rotenone Sigma R8875, Antimycin A Sigma A8674). For the glycolytic stress test, we used Glucose (100mM, 20μL port A; Sigma G8270), Oligomycin (20μM, 22μL port B; Sigma 75351), and 2-Deoxyglucose (500mM, 25μL port C; Sigma D6134). Quantification of the parameters generated in each test were performed using the formulas provided by Agilent. Data represented as ECAR/OCR fold change were normalized to the average ECAR/OCR of control unstimulated BMDMs.

### LiCAT-seq

Low-input chromatin accessibility and transcriptome sequencing (LiCAT-seq) method was adapted from (Liu *et al*., 2019). To separate the RNA containing cytosol from the nucleus, sorted cells were lysed in 6.5uL of mild lysis buffer (10mM NaCl, 10mM Tris-HCL, pH 7.5; 0.5% NP-40, 0.25uL of 40 U/uL RNase-inhibitor [NEB M0314] and nuclease free water) for 30’ at 4^oC^. The lysis was vortexed for 1’ prior to be centrifuged at 2000 g for 5’ at 4^oC^. The top 4uL of lysis supernatant containing the RNA fraction was directly placed into 350uL of RLT buffer from an RNEasy micro kit (Qiagen 74004), and RNA was purified the following manufacturer’s instructions. The remaining lysis supernatant containing the nuclei were immediately placed into a transposase reaction using the Tagment DNA TDE1 enzyme kit (Illumina 20034197) according to manufacturer’s instructions, and placed in a 37^C^ shaker for 30’. DNA was immediately purified using the DNA Clean and Concentrator-5 kit (Zymo Research D4013) according to manufacturer’s instructions.

### ATAC-seq analysis

The quality of the fastq files was accessed using FastQC (Simon Andrews, 2020). Illumina adapters and low-quality reads were filtered out using BBDuk (http://jgi.doe.gov/data-and-tools/bb-tools). Bowtie2 (v.2.3.4.3) (Langmead and Salzberg, 2012) was used to align the sequencing reads to the mm10 reference murine genome. Samtools (v.1.11) (H. Li *et al*., 2009) was used to select the mapped reads (samtools view -b -q 30) and sort the bam files. PCR duplicates were removed using Picard MarkDuplicates tool (Broad Institute, 2009). The normalization ratio for each sample was calculated by dividing the number of uniquely mapped murine reads of the sample with the lowest number of reads by the number of uniquely mapped murine reads of each sample. These normalizations ratios were used to randomly sub-sample reads to obtain the same number of reads for each sample using using samtools view -s. Bedtools genomecov was used to create bedgraph files from the bam files (Quinlan and Hall, 2010). Bigwig files were created using deepTools bamCoverage (Ramírez *et al*., 2016). Peaks were called using MACS2 (v2.1.2) (Zhang *et al*., 2008) with ENCODE recommended parameters. IDR was used to identify the reproducible peaks between the replicates (Li *et al*., 2011). Further processing of the peak data was performed in R. Average profiles and heatmaps were generated using ngs.plot (Shen *et al*., 2014).

### Gene expression analysis

RNA-sequencing analysis was performed on RNA obtained from three independent biological replicate pools of double sorted 5,000 HSC, 10,000 MPP3, 50,000 GMP, and 50,000 BM Mon. RNA was obtained using the Qiagen RNAEasy Micro kit (Qiagen 74004) according to manufacturer’s instructions immediately after cell sorting. RNA quality was verified using a High Sensitivity ScreenTape Assay on the Tape Station 2200 (Agilent Technologies) and measured with a NanoDrop 1000 (Thermo Fisher Scientific). Library construction was performed using the Universal Plus mRNA Library Kit (NuGen Technologies), and sequencing was performed on the NovaSeq 6000 instrument (Illumina) using paired-end sequencing (2 3 150 bp) by the University of Colorado Cancer Center Genomics and Microarray Core. Illumina adapters were trimmed using BBDuk (Cite BBTools) and reads <50bp post trimming were discarded. Reads were aligned and quantified using STAR (2.6.0a) (Dobin *et al*., 2013) against the Ensembl mouse transcriptome (GRCm38.p6 genome (release 96)). Reads were normalized to counts per million (CPM) using the edgeR R package (Robinson, McCarthy and Smyth, 2009). Differential expression for the indicated comparisons were calculated using the limma R package and the voom function (Ritchie *et al*., 2015). Over-representation analysis was performed with clusterProfiler R (Yu *et al*., 2012) and gene sets from the Molecular Signatures Database (Liberzon *et al*., 2011) with a gene threshold of FDR <0.5.

### Metabolomics analysis

300,000 BMDMs were sorted and metabolomics analyses were performed via ultra-high pressure-liquid chromatography-mass spectrometry (UHPLC-MS – Vanquish and Q Exactive, Thermo Fisher) as previously reported (Nemkov, D’Alessandro and Hansen, 2015). Briefly, cells were extracted in ice cold methanol:acetonitrile:water (5:3:2 v/v/v) at a concentration of 2 million cells/mL of buffer. After vortexing for 30 min at 4°C, samples were centrifuged at 12,000 g for 10 min at 4°C and supernatants processed for metabolomics analyses. Ten microliters of sample extracts were loaded onto a Kinetex XB-C18 column (150 × 2.1 mm i.d., 1.7 μm – Phenomenex). A 5 min gradient (5%–95% B, phase A: water + 0.1% formic acid and phase B: acetonitrile with + 0.1% formic acid for positive ion mode; 0%–100% B, phase A: 5% acetonitrile + 5mM ammonium acetate and phase B: 95% acetonitrile + 5mM ammonium acetate for negative ion mode) were used to elute metabolites. The mass spectrometer scanned in Full MS mode at 70,000 resolution in the 65–975 m/z range, 4 kV spray voltage, 45 sheath gas and 15 auxiliary gas, operated in negative and then positive ion mode (separate runs). Metabolite assignment was performed against an in-house standard library, as reported (Nemkov, D’Alessandro and Hansen, 2015; Nemkov *et al*., 2019).

### Statistical analysis

Statistical analysis was performed using Prism 9 software (Graphpad). *P* values were determined using a Mann-Whiteny *U*-test, one-way analysis of variance, or two-way analysis of variance. *P*-values ≤ 0.05 were considered statistically significant.

## RESULTS

### Chronic autoimmune inflammation potentiates myelopoiesis

To understand how autoimmune inflammation impacts hematopoiesis and HSC function, we used the pristane-induced mouse model of the AD systemic lupus erythematosus (SLE). The pristane model is the most widely accepted model of environmentally induced SLE, and faithfully recapitulates many aspects of human SLE disease, including production of anti-nuclear autoantibodies, a type-I interferon signature (IFN-I), and kidney nephritis (Satoh and Reeves, 1994; Reeves *et al*., 2009; Freitas, de Oliveira and Monticielo, 2017). To investigate the impact of chronic autoimmune inflammation on the hematopoietic system, (**Fig 1A**), we analyzed hematopoietic populations in the peripheral blood (PB) and bone marrow (BM) of mice treated ± pristane for 8 weeks. All mice treated with pristane had overt inflammatory myelopoiesis evidenced by a significant increase in PB neutrophils (**Fig 1B**) and granulocytes (Gr) (**Fig 1C**) as determined by CBC and flow cytometry, respectively. PB lymphocytes from pristane-treated mice also had increased surface expression of the interferon sensitive gene (ISG) Sca-1 on PB T and B cells (**Fig 1D, S1A**), a validated marker of SLE progression (Xu *et al*., 2014). Pristane treatment did not alter BM cellularity (**Fig 1E**). However, pristane-treated mice had a significant increase in the number of pre-Gr and Gr in the BM (**Fig 1F**), with a concomitant decrease in BM B and T lymphocytes. (**Fig S1B-D**). Overall, these data are consistent with preferential generation of mature granulocytes at the expense of the lymphoid lineage, indicative of inflammatory myelopoiesis (King and Goodell, 2011; Caiado, Pietras and Manz, 2021).

**Figure 1.**
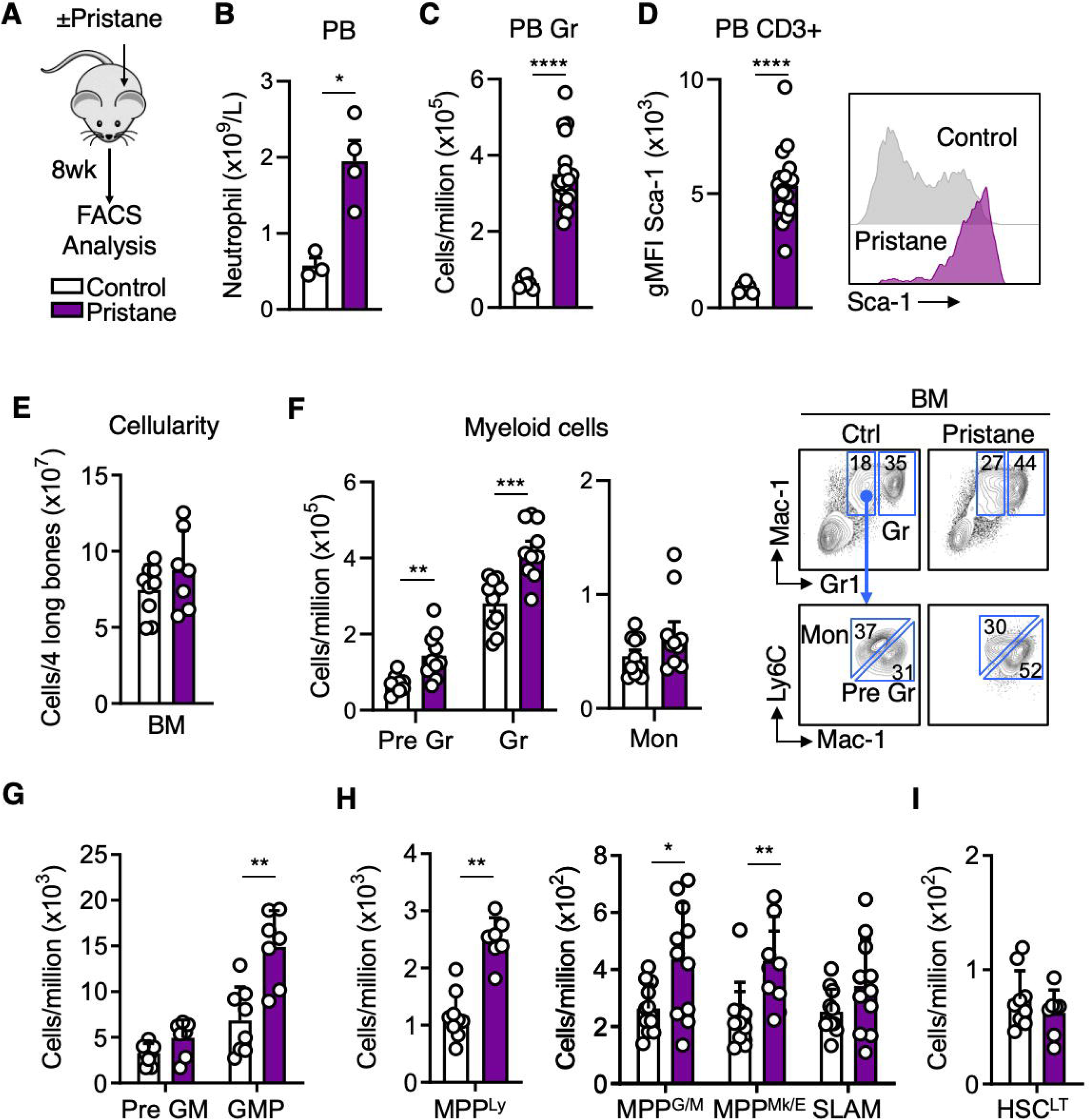
Chronic autoimmune inflammation potentiates myelopoiesis. **(A-I)** Experimental Design; mice were treated ± pristane for eight weeks for FACS analysis. **(B)** Complete blood count (CBC) analysis of peripheral blood (PB) neutrophils; n = 3 Control, 4 Pristane. **(C)** Quantification of PB granulocytes via flow cytometry; n = 6 Control, 18 Pristane. **(D)** Flow cytometry analysis of geometric mean fluorescent intensity (gMFI) of Sca-1 on PB CD3+ lymphocytes (left), and representative histograms (right); n = 6 Control, 18 Pristane. **(E)** Enumeration of bone marrow (BM) cellularity in the four long bones of the legs; n = 9 Control, 7 Pristane. **(F-I)** Quantification of FACS analysis of (F) mature myeloid cells (left) and representative FACS plots (right), (G) myeloid progenitor cells, (H) immature hematopoietic progenitors, and (I) long-term HSC (HSC^LT^); n = 7-11/group See figure S2 for surface marker definitions and representative FACS plots of indicated populations. Data are represented as SEM. *p < 0.05, **p < 0.01, ***p < 0.001, ****p < 0.0001, significant versus control; significance determined by two-tailed Mann Whitney U-test.

We next assessed whether increased myeloid output was accompanied by changes in HSPC populations. Because the pristane mouse model induces potent release of type-I interferons (IFN-I) (Freitas, de Oliveira and Monticielo, 2017), traditional flow cytometry gating strategies cannot be used to accurately interrogate HSPC due to increased Sca-1 expression in HSPC that do not ordinarily express this marker (Pietras *et al*., 2014). Instead, we substituted Sca-1 with expression of the surface marker ESAM to delineate between myeloid progenitors (traditionally Lin^-^/Sca-1^-^/cKit^+^; herein Lin^-^/ESAM^-^/cKit^+^) and immature HSPCs with multilineage potential (traditionally Lin^-^/Sca-1^+^/cKit^+^; herein Lin^-^/ESAM^+^/cKit^+^; LEK) (Camilla Forsberg *et al*., 2005; Ooi *et al*., 2009; Pietras *et al*., 2014) (**Fig S2A, B**). Indeed, we found that Sca-1 expression was significantly increased on all HSPC populations, particularly GMPs (**Fig S1F**), emphasizing the requirement for using ESAM to accurately enumerate myeloid progenitors in this setting. Consistent with elevated myeloid cell abundance in the PB and BM, the number of granulocyte-monocyte progenitors (GMP) was significantly increased in pristane-treated mice (**Fig 1G**), with a corresponding decrease in the number of common lymphoid progenitors (CLP; **Fig S1E**). Interestingly, all three lineage-primed multipotent progenitor (MPP) populations MPP^Ly^ (formerly MPP4), MPP^G/M^ (formerly MPP3), and MPP^Mk/E^ (formerly MPP2) were significantly expanded, whereas the phenotypic HSC-enriched SLAM population (LEK/Sca1^+^/Flk2^-^/CD48^-^/CD150^+^; SLAM) and the proportion of EPCR^+^/CD34^-^ long-term HSC (HSC^LT^) (Balazs *et al*., 2006; Wilson *et al*., 2008; Rabe *et al*., 2020) therein were unchanged (**Fig 1H, I**). Further, we found no change in cell cycle activity in SLAM after eight weeks of pristane treatment (**Fig S1G**). Overall, these data show that the autoimmune inflammation caused by pristane treatment induces myeloid expansion in the BM and peripheral blood.

### Chronic autoimmune inflammation activates defense programs in HSPC and monocytes

To further understand molecular mechanisms associated with the expansion of the myeloid lineage, we performed RNA sequencing (RNA-seq) on cell populations along the entire myeloid hierarchy including BM monocytes (BM Mon), GMP, the myeloid biased MPP^G/M^, and the HSC-enriched SLAM fraction (SLAM) (**Fig 2A**). We investigated the transcriptome at four weeks post pristane treatment, to better understand the gene expression changes associated with the phenotypic alterations in the blood system at eight weeks. First, we ran unsupervised analysis of all cell types using principal component analysis (PCA). Notably, HSPC clustered primarily based on their identity rather than due to treatment, validating the phenotypic definitions we used to prospectively isolate them. Only BM Mon completely segregated between control and pristane treated conditions, though they remained closely associated in PCA space (**Fig 2B**). Using an adjusted p-value (p-adj) of 0.05 as a cutoff, we identified 3,044 differentially expressed genes (DEG) in Mon, 2,358 in GMP, 469 in MPP^G/M^, and 231 DEG in the SLAM fraction (**Fig 2C, Table S1**). We next ran overrepresentation analysis (ORA) on the DEGs in each population. Interestingly, GMP and BM Mon were significantly enriched for activated gene programs related to metabolism and cellular activation, including glycolytic and oxidative metabolism, antigen presentation via MHC-I, and activation of Myc and mTOR (**Fig 2D**). In comparison, MPP^G/M^ and SLAM were significantly enriched for multiple transcriptional programs related to interferon signaling and immune effector processes (**Fig 2E**). There were also cell-specific gene expression changes associated with pristane treatment, with BM Mon further activating innate immunity programs and GMP potently engaging ribosome and mitochondrial biogenesis pathways (**Fig S3A**). Interestingly, MPP^G/M^ activated gene programs related to cellular differentiation and translation, while SLAM activated innate immune defense programs (**Fig S3B**). Conversely, BM Mon and GMP were enriched for downregulated genes related to cell motility and wound healing responses (**Fig S3C**) whereas MPP^G/M^ and SLAM downregulated pathways related to kinase signaling activity (**Fig S3D**). Together, these data show that pristane-mediated inflammation triggers broad activation of inflammatory and host defense genes in HSPC and monocytes, whereas induction of metabolic genes closely associates with myeloid lineage commitment.

**Figure 2.**
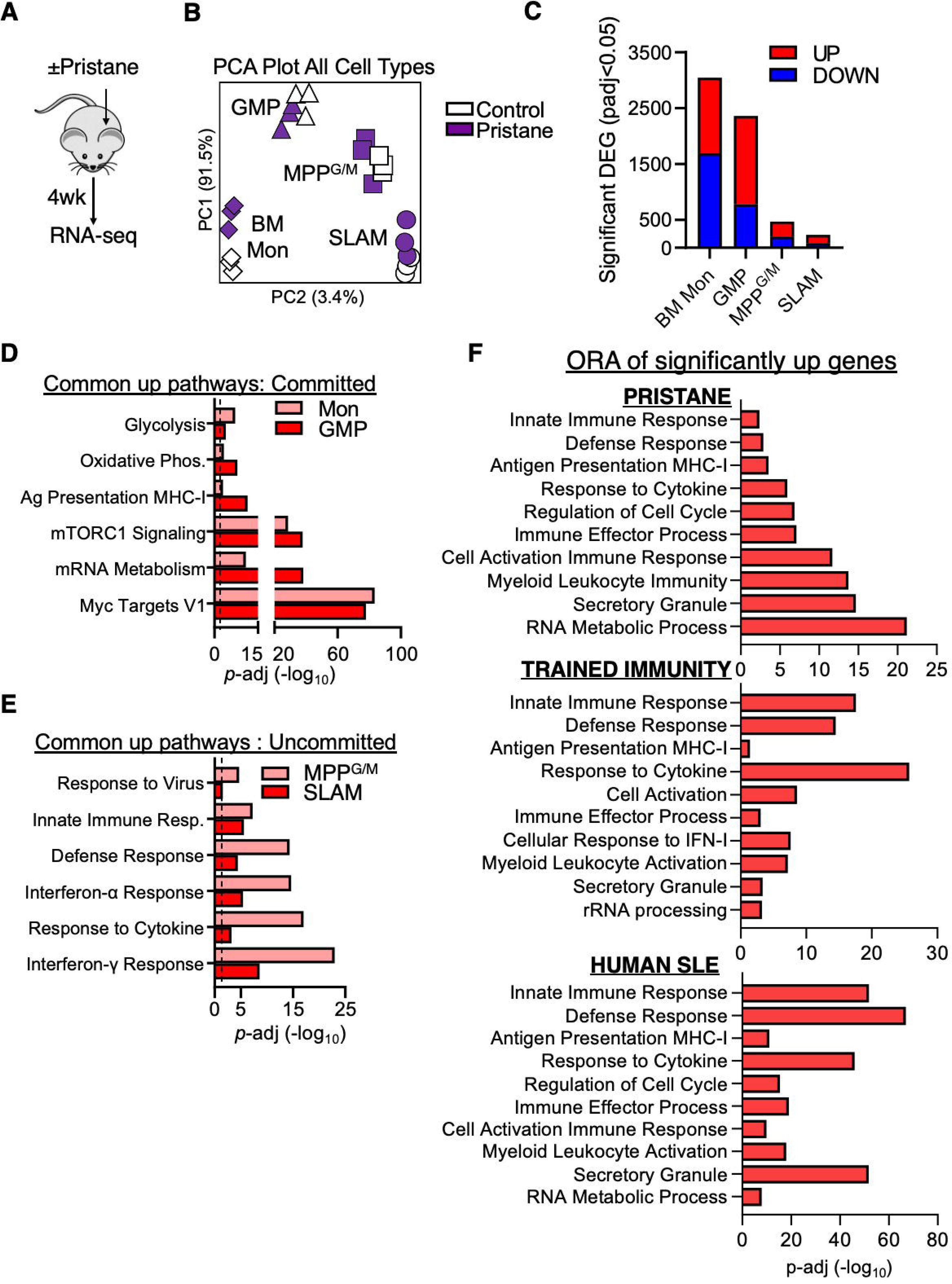
Pristane treatment activates defense programs in HSPC and monocytes. **(A-E)** Experimental Design; mice were treated ± pristane for four weeks for RNA-seq analysis; n = 3/group **(B)** Principal component analysis (PCA) of bone marrow monocytes (BM Mon), granulocyte monocyte progenitors (GMP), myeloid biased multipotent progenitors (MPP^G/M^), and the HSC enriched SLAM fraction (SLAM). **(C)** Quantification of significant differentially expressed genes (DEG) with an adjusted p value (padj) < 0.05. **(D-E)** Over-representation analysis (ORA) of significantly enriched pathways (p-adj < 0.05) determined using the significantly up-regulated DEG (From C) in (D) BM Mon and GMP, or (E) MPP^G/M^ and SLAM; significance cut-off of p-adj < 0.05 is represented with a dashed line. **(F)** ORA analysis of significantly enriched pathways (p-adj < 0.05) determined using significantly up-regulated DEG (p-adj < 0.05) as determined by RNA-seq analysis: of BM Mon from pristane treated mice (From C) (Top), of human PBMCs from models of TI(middle), and of human PBMCs from SLE patients (bottom).

We noted that the defense response, interferon, and metabolic programs in BM Mon from pristane-treated mice resembled gene expression patterns associated with trained immunity (TI) as well as myeloid cells from autoimmune disease patients (Kaufmann *et al*., 2018; Mitroulis *et al*., 2018; Rönnblom and Leonard, 2019). To address the extent to which these signatures overlap with our RNA-seq datasets, we first compiled gene lists of upregulated DEG from published data sets of human PBMCs trained with β-glucan, BCG vaccination, and malaria infection (Novakovic *et al*., 2016, GSE85243; Moorlag, Khan, *et al*., 2020, GSE141656; Walk *et al*., 2020, GSE137044) (2,404 genes) (**Table S2**), or SLE PBMCs (Nakou *et al*., 2008; Smiljanovic *et al*., 2012; Kennedy *et al*., 2015 GSE50772; Labonte *et al*., 2018; Grigoriou *et al*., 2020) (3,648 genes) (**Table S3**). The results from all three analysis show a high degree of similarity, with overlapping activation of innate immune response, interferon response, and antigen presentation gene programs (**Fig 2F**). These results indicate that the pristane model induces gene programs associated with human AD and immune training and suggest the three settings may share a common biology.

### BMDMs from autoimmune mice have increased inflammatory function

Given the link between our RNA-seq results, human AD, and immune training datasets, we next assessed whether chronic autoimmune inflammation enhances myeloid cell function in a manner consistent with immune training. The two major functional hallmarks of trained immunity are increased pathogen killing activity and inflammatory cytokine production in response to a subsequent immune challenge (Bekkering *et al*., 2021). Hence, we used bone marrow-derived macrophages (BMDMs) as a well-characterized system for studying the functional and mechanistic features of immune training (van’t WouT, Poell and van Furth, 1992; Kaufmann *et al*., 2018). We generated BMDMs from mice treated ± pristane for eight weeks, and as controls we included BMDMs from mice treated for seven days with poly I:C (pIC), a double-stranded RNA analogue which induces IFN-I driven inflammation independently of an autoimmune process (Pichlmair and Reis e Sousa, 2007; Essers *et al*., 2009; Pietras *et al*., 2014). We then confirmed the generation of a highly pure BMDM population in each setting (**Fig S4A**). To assess the extent of immune training, we used a live killing assay in which BMDMs were challenged with the intracellular bacterium *Mycobacterium avium* (*M. avium*) for 72hrs (**Fig 3A**). We chose *Mycobacterium avium* as our challenge because it serves as an experimental model of M. tuberculosis infection, a pathogen immune training is known to increases protection against (Gutierrez *et al*., 2004; Kaufmann *et al*., 2018; Moorlag, Khan, *et al*., 2020; Zhou *et al*., 2021). Interestingly, BMDMs from mice treated with pristane or pIC had an increased ability to kill live *M. avium* based on qRT-PCR analysis of bacterial copy number after 72 hours (**Fig 3B**). We also found that after 4 hours of co-culture with *M. avium*, the BMDMs from pristane and pIC treated mice had a significant increase in the expression of Ly6C, which marks phagocytes with elevated antimicrobial activity, compared to BMDMs from control mice (**Fig 3C**) (Serbina *et al*., 2003; Pietras *et al*., 2011). We also harvested culture supernatant from the killing assay to investigate levels of inflammatory cytokines produced by the BMDMs. Notably, there was a significant increase in the amount of TNFα, and a trending increase in IL-6, produced by BMDMs from pristane-treated mice compared to control mice (**Fig 3D, S3E**). Collectively, these data show that BMDMs from pristane treated mice exhibit hallmark features of trained immunity. Interestingly, seven days of pIC treatment was sufficient to recapitulate key aspects of this phenotype, consistent with evidence for immune training in response to even a single pIC injection in mice (de Laval *et al*., 2020)

**Figure 3.**
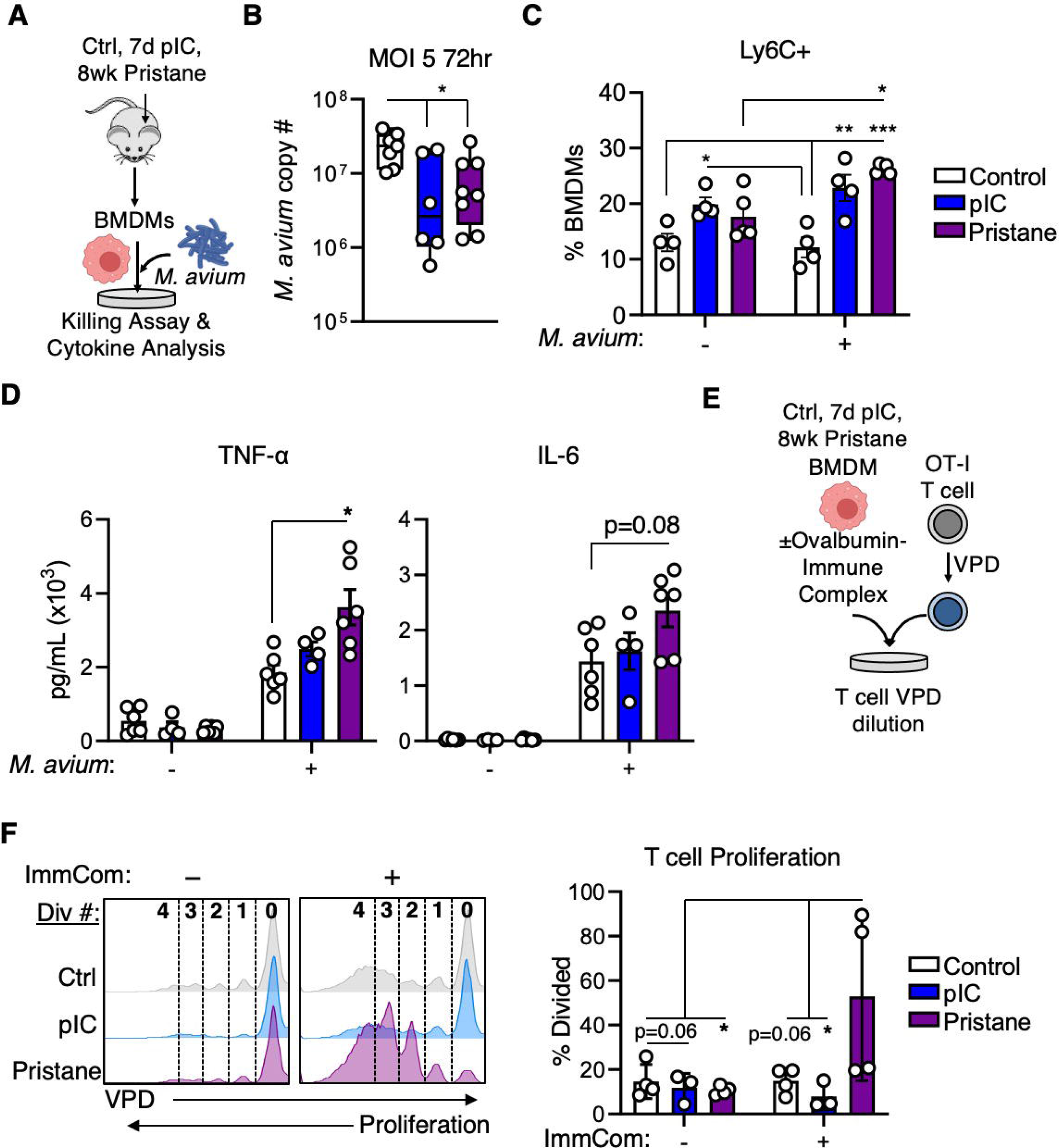
BMDMs from autoimmune mice have increased inflammatory function. **(A-D)** Experimental design; mice were treated ± pristane for eight weeks or pIC for seven days, then bone marrow derived macrophages (BMDMs) were generated and placed into a killing assay co-culture with *Mycobacterium avium* for 72 hours before assessing bacterial load, and cytokines generated in the culture supernatant. **(B)** Quantification of *Mycobacterium avium* (*M. avium*) copy number using qRT-PCR from co-culture with BMDMs after 72 hours with a multiplicity of infection (MOI) dose of 5; n = 7 Control, 6 pIC, 8 Pristane. **(C)** Percent of BMDMs that gained surface expression of Ly6C after four hours of culture with *M. avium* (From B); n = 4/group **(D)** Cytokine analysis of the culture supernatant from the BMDM and *M*. *avium* killing assay after 72 hours (From B) of TNF-α (left) and IL-6 (right); n = 6 Control, 4 pIC, 6 Pristane. **(E-F)** Experimental design; BMDMs were generated from mice that were treated ± pristane for eight weeks or pIC for seven days, and were then stimulated overnight with immune complexes (ImmCom) of IgG targeting Ovalbumin protein, before being placed in a co-culture assay with violet proliferative dye (VPD) labelled OT-I T cells for three days. **(F)** Representative histograms (left) of VPD dilution in T cells after three days of culture with ImmCom stimulated BMDMs (From E) and quantification of the percent of T cells that divided in the culture (right); n = 4 Control, 3 pIC, 4 Pristane. Data are represented as SEM. *p < 0.05, **p < 0.01, ***p < 0.001, significance determined by One-way ANOVA with Tukey’s post-test (B, D), or Two-way ANOVA with Tukey’s post-test (C, F)

Bystander activation of T cells by macrophages is postulated to play a role in activation of the autoimmune cascade (Fujinami *et al*., 2006; Pacheco *et al*., 2019). Thus, we tested the capacity of BMDMs from pIC- or pristane-treated mice to activate OT-I antigen specific T cells. As immune complex deposition is a key driver of inflammation and SLE pathogenesis in the kidneys and other tissues, we chose to restimulate our BMDMs with *in vitro* generated immune complexes (ImmCom; complexes of antibodies targeting their antigen, herein IgG targeting Ovalbumin (OVA) protein) as a physiologically relevant stimulus (Satoh *et al*., 1995; Herrada *et al*., 2019). After stimulating our BMDMs with the OVA-ImmCom overnight, we co-cultured them with violet proliferative dye (VPD)-labeled OTI-T cells for three days (**Fig 3E**). After the incubation period the T cells were isolated, and their proliferative activity quantified by VPD dilution. Strikingly, BMDMs from pristane-treated mice robustly induced T cell proliferation compared to stimulated BMDMs from control or pIC treated mice, based on the total number of T cells that have divided (**Fig 3F**). Hence, our data show that exposure to chronic autoimmune inflammation potentiates the capacity of myeloid cells to present antigens to T cells. These findings complement, from a functional standpoint, the upregulation of antigen presentation gene signatures in monocytes from pristane-treated mice.

### Pristane exposure triggers glycolytic metabolism in bone marrow-derived macrophages

The increased inflammatory response characteristic of trained immunity in macrophages has been mechanistically linked to increased glycolytic metabolism (Cheng *et al*., 2014; Arts *et al*., 2016). Given BMDMs from pristane-treated mice exhibited functional properties consistent with immune training, we next assessed whether changes in their metabolic properties were consistent with a trained phenotype. Thus, we again generated BMDMs from mice treated ± pristane for eight weeks or with pIC for seven days and stimulated them overnight with ImmCom (IgG specific for NP-BSA_16_) (**Fig 4A**). We then performed glycolytic and mitochondrial stress tests using the Seahorse extracellular flux assay. Consistent with prior descriptions of metabolic reprogramming associated with trained immunity, we found that BMDMs from pIC- and pristane-treated mice had a significant increase in glycolysis, as well as glycolytic capacity (a measure of the maximum rate of glycolysis that a cell can achieve) compared to BMDMs from control mice following ImmCom stimulation (**Fig 4B, C**). Interestingly, only BMDMs from pristane-treated mice had a significant increase in glycolytic reserve (the change in ECAR between glycolysis and glycolytic capacity, indicative of capacity to engage a stress response) following ImmCom stimulation (**Fig. 4C**), perhaps reflective of the duration of inflammation versus pIC. Likewise, we found that BMDMs from pristane-treated mice show increased oxidative phosphorylation (OXPHOS) activity, including a significant increase in basal respiration, maximum respiration, and spare capacity (the difference between basal and maximum respiration) compared to BMDMs from control mice (**Fig. S4B, C**). Interestingly, the rates of OXPHOS did not change in response to ImmCom stimulation, consistent with a reported shift of macrophages towards glycolysis when stimulated with ImmCom (Jing *et al*., 2020). These data show that BMDMs derived from pristane-treated mice retain an activated glycolytic profile that can be triggered by an immune stimulus.

**Figure 4.**
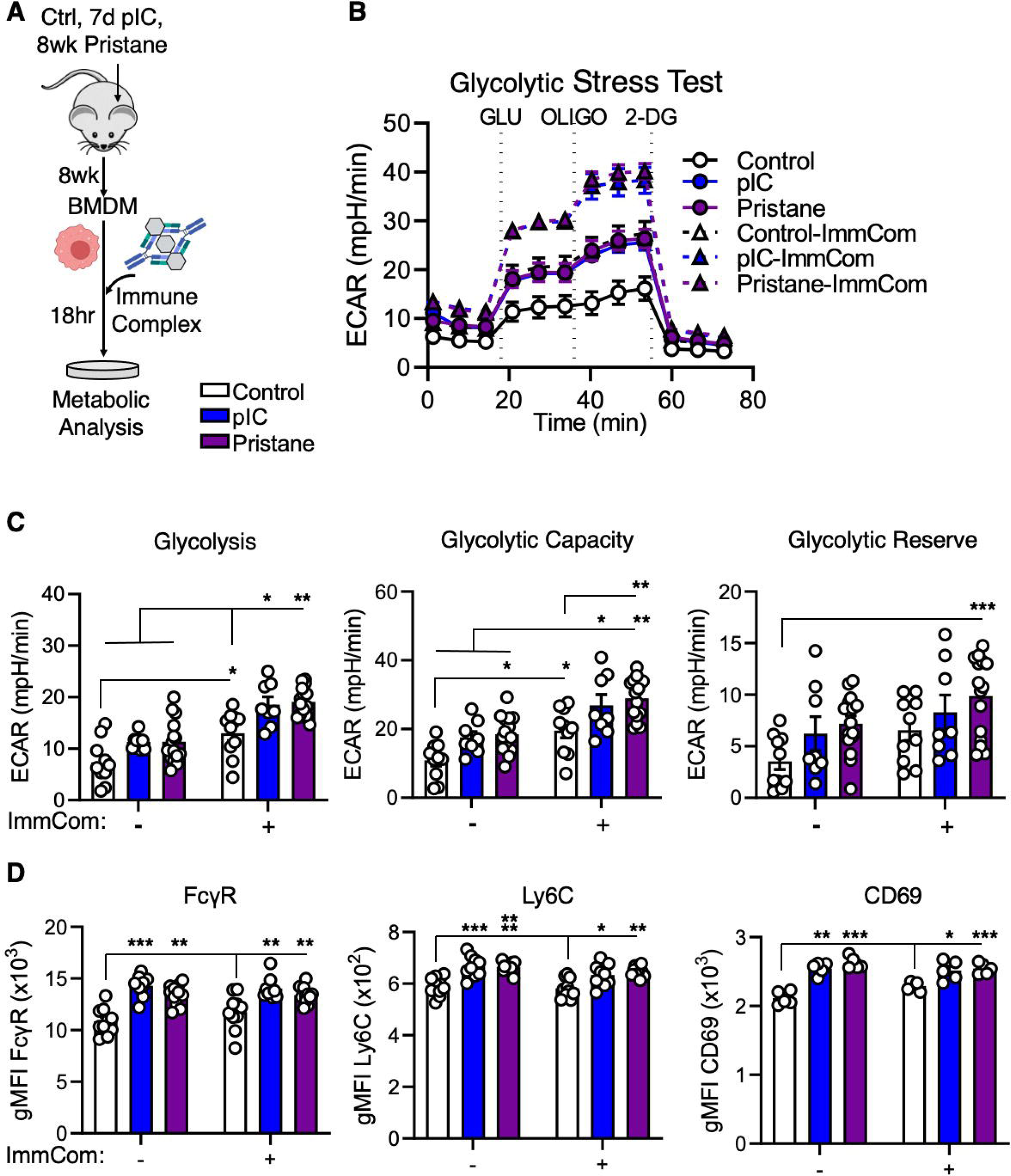
Pristane exposure triggers glycolytic metabolism in BMDMs. **(A-C)** Experimental design; bone marrow derived macrophages (BMDMs) were generated from mice that were treated ± pristane for eight weeks or pIC for seven days, and were then stimulated overnight with immune complexes (ImmCom) of IgG targeting NP-BSA_16_ before having their glycolytic metabolism assessed using the Seahorse glycolytic stress test. **(B)** Tracer of glycolytic stress test of unstimulated (solid lines) and ImmCom stimulated (dashed lines) BMDMs. n = 14 Control, 8 pIC, 15 Pristane. **(C)** Quantification of glycolysis (ECAR after the addition of glucose (GLU); left), glycolytic capacity (ECAR after the addition of oligomycin (OLIGO), middle), and glycolytic reserve (the difference in ECAR between glycolysis and glycolytic capacity, right); n = 14 Control, 8 pIC, 15 Pristane. **(D)** BMDMs were generated from mice that were treated ± pristane for eight weeks or pIC for seven days, and then stimulated with ImmCom prior to quantifying the surface expression of the indicated proteins using geometric mean fluorescence intensity (gMFI) FACS analysis. BMDMs were either stimulated with ImmCom for 24hr (FcγR left, and Ly6C middle) or 72hr (CD69 right); CD69 gMFI data are representative of two independent experiments; n = 10 Control, 9 pIC, 11 Pristane. Data are represented as SEM. *p < 0.05, **p < 0.01, ***p < 0.001, ****p < 0.0001, significance determined by Two-way ANOVA with Tukey’s post-test.

Given the increased inflammatory and metabolic activity in BMDMs from pristane treated mice, we analyzed expression of key surface markers by flow cytometry. Consistent with increased metabolic and functional activity in response to ImmCom stimulation, we found that BMDMs from mice treated with pIC or pristane had significantly increased surface expression of FcγR (**Fig. 4D, S4D**), which is consistent with analysis of PB monocytes in SLE patients (Li *et al*., 2009; Li *et al*., 2010; Abd-Elhamid *et al*., 2017). In line with their pro-inflammatory phenotype, these BMDMs also exhibited increased levels of Ly6C as well as CD69, a type II C-lectin receptor that can facilitate glucose uptake and mTOR activity, leading to activation of nitric oxide and TNF-α (Conde *et al*., 1996; Marzio *et al*., 1997; Cibrián and Sánchez-Madrid, 2017). Of note, the increased surface expression of these proteins was evident even in unstimulated cells, perhaps reflective of central immune training in the mouse BM. Overall, these data show that BMDMs from pristane and pIC-treated mice exhibit metabolic and phenotypic characteristics consistent withTI.

### HSC^LT^ from autoimmune mice are a transplantable reservoir for trained macrophages

As myeloid cells are relatively short-lived, the HSPC pool has been speculated to serve as a ‘memory’ compartment for TI, and it has been shown that HSPC stimulated with LPS or β-glucan can generate trained macrophages following transplantation into recipient mice (Kaufmann *et al*., 2018; de Laval *et al*., 2020). Given BMDMs from pristane-treated mice exhibit a trained immunity phenotype, we wanted to investigate whether HSC function as a long-term reservoir for trained immunity in response to chronic autoimmune inflammation. To ensure rigorous prospective isolation of functional HSC in the setting of inflammation, we transplanted purified LEK/Sca1^+^/CD48^-^/CD150^+^/ EPCR^+^/CD34^-^ HSC^LT^ cells from mice treated ± pristane for eight weeks or pIC for seven days into lethally irradiated recipient mice (**Fig 5A**). Our previous studies have shown that the EPCR^+^/CD34^-^ SLAM cell fraction enriches for cells that retain long-term stem cell activity even in the presence of inflammation (Rabe *et al*., 2020). Post-transplant, all recipient mice had equivalent levels of donor chimerism in the PB regardless of the HSC^LT^ donor without any evidence of donor lineage skewing (**Fig 5B, S5A, B**). Further, after 18 weeks we found that all recipient mice achieved high levels of BM chimerism (**Fig S5C**). When we investigated the donor chimerism within specific BM hematopoietic cell populations, we found equivalent donor chimerism in mature myeloid (**Fig S5D**) and lymphoid fractions (**Fig S5E**), as well as GMP, MPP^LY^, MPP^G/M^, MPP^Mk/E^, and SLAM (**Fig S5F, G**), with a trend towards reduced HSC^LT^ chimerism, perhaps indicative of a minor reduction in repopulating capacity following chronic inflammatory stress. We did not observe evidence of increased autoimmune inflammatory activity in BM HSPC based on Sca-1 levels, indicating that we did not transplant pristane-induced disease (**Fig S5H, I**). Of note, these data contrast with our previous study showing reduced repopulating activity of HSC from pIC-treated mice (Pietras *et al*., 2014), which were based on the less-stringent SLAM definition that does not exclude EPCR-cells with extensive short-term Mk activity (Haas *et al*., 2015; Chavez *et al*., 2022). Taken together, these data indicate HSC^LT^ retain long-term multilineage repopulating activity in the setting of chronic autoimmune inflammation.

**Figure 5.**
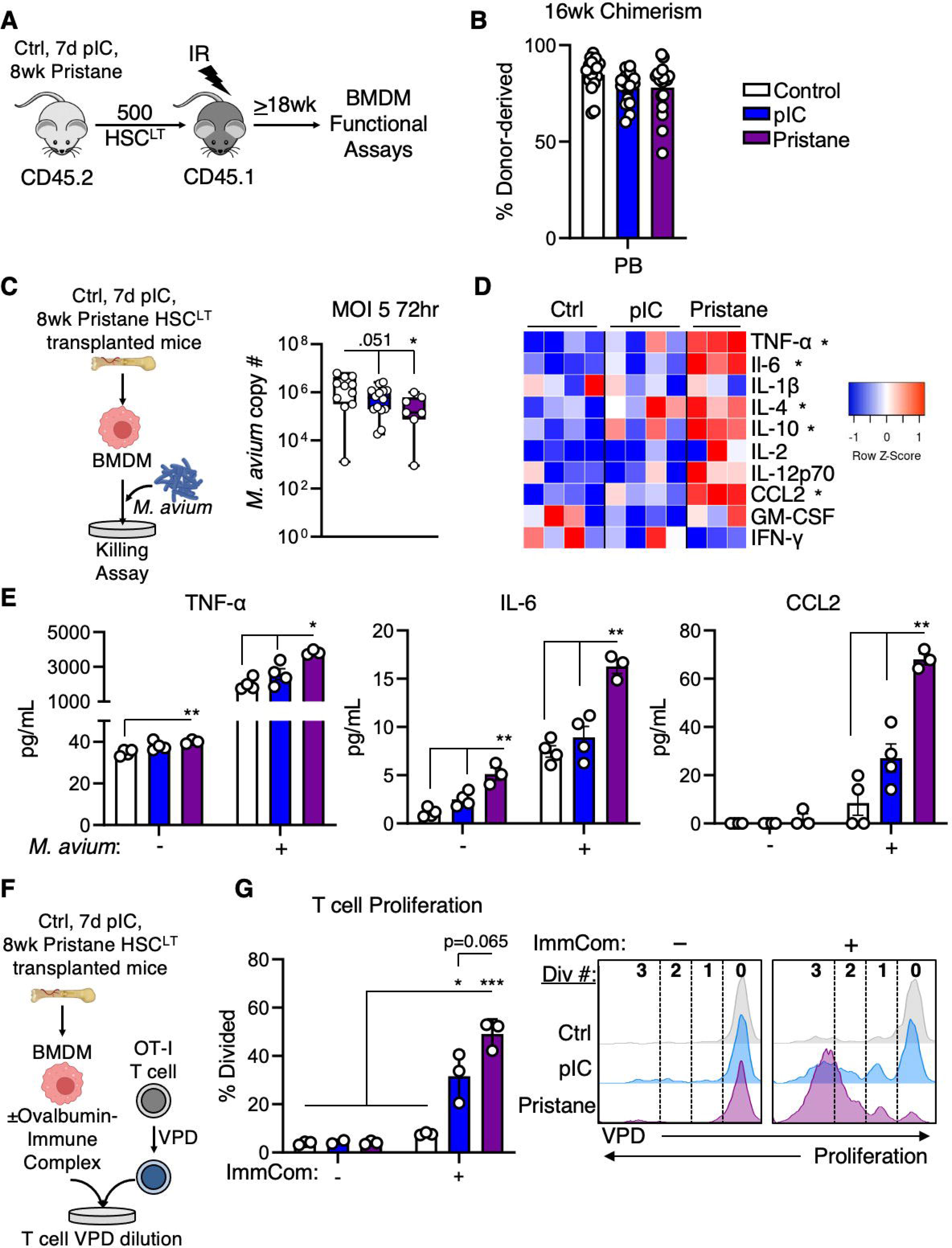
HSC^LT^ from autoimmune mice are a long-term reservoir for trained BMDMs. **(A-G)** Experimental design; mice were treated ± pristane for eight weeks or pIC for seven days, then 500 HSC^LT^ (LEK/Sca1^+^/Flk2^-^/CD48^-^/CD150^+^/EPCR^+^/CD34^-^) were FACS sorted from CD45.2 donor mice and then transplanted into lethally irradiated CD45.1 recipient mice. **(B)** Percent of donor chimerism in the peripheral blood of mice transplanted with HSC^LT^ from control, pIC-, or pristane-treated donor mice (From A) after 16 weeks of engraftment; n = 17 Control, 16 pIC, 19 Pristane. **(C)** Experimental design (left); bone marrow derived macrophages (BMDMs) were generated from mice transplanted with HSC^LT^ from control, pIC-, or pristane-treated donor mice (From A), and were then placed into a killing assay co-culture with *Mycobacterium avium* (*M. avium*) for 72 hours, and quantification of *M. avium* copy number using qRT-PCR (right); n = 11 Control, 14 pIC, 7 Pristane. **(D-E)** Heatmap of cytokines (D) and quantification (E) of TNF-α (left), IL-6 (middle), and CCL2 (right) from the cell culture supernatant at the end of the 72hr killing assay (From C); (D) significance to Control; n = 4 Control, 4 pIC, 3 Pristane. **(F-G)** Experimental design (F); bone marrow derived macrophages (BMDMs) were generated from mice transplanted with HSC^LT^ from control, pIC-, or pristane-treated donor mice (From A), and were then stimulated overnight with immune complexes (ImmCom) of IgG targeting Ovalbumin protein before being placed in a co-culture assay with violet proliferative dye (VPD) labelled OT-I T cells for three days. (G) Percent of T cells that divided after the three-day co-culture (left), and representative histograms of VPD dilution (right); n = 3/group. Data are represented as SEM. *p < 0.05, **p < 0.01, ***p < 0.001, significance determined by Two-way ANOVA with Tukey’s post-test.

To assess whether BMDMs derived from donor HSC^LT^ from pristane-treated mice retained a trained phenotype, we generated BMDMs from our recipient mice and tested them in the same functional assays as before. Consistent with a trained phenotype, BMDMs derived from pristane- and pIC-exposed donor HSC^LT^ retained enhanced bacterial killing capacity (**Fig 5C**), as evidenced by significantly lower *M. avium* copy numbers compared to BMDMs from mice transplanted with control HSC^LT^. We also assessed the levels of pro-inflammatory cytokines present in the infection co-culture supernatant. We found that BMDMs derived from pristane-exposed donor HSC^LT^ produced higher levels of TNF-α, IL-6, and the myeloid chemokine CCL2, as well as IL-10 and IL-4, though this phenotype was not shared by BMDMs from pIC-exposed donor HSC^LT^ (**Fig 5D, E**). Importantly, we found that BMDMs from pristane- and pIC-exposed donor HSC^LT^ retained the capacity to stimulate OT-I T cells to proliferate following ImmCom stimulation (**Fig 5F, G**). Collectively, these data demonstrate that HSC^LT^ give rise to trained macrophages, even after transplantation.

### Pristane exposure induces heritable molecular reprogramming of HSC^LT-^derived GMP

To address whether pristane exposure induces heritable molecular changes in HSC^LT^-derived myeloid progenitor cells, we investigated transcriptional and epigenetic changes underlying the inherited program. Thus, we isolated donor-derived GMP from our transplanted mice and performed low-input chromatin accessibility and transcriptomics sequencing (LiCAT-seq) (Liu *et al*., 2019) to obtain RNA-seq and ATAC-seq libraries from the same pools of cells (**Fig 6A**). Previous literature has shown that HSPC exhibit an increase in chromatin accessibility at inflammatory genes following immune stimuli in primary mice, which may not represent a true HSC-inherited phenotype (de Laval *et al*., 2020). Changes in chromatin accessibility have not to our knowledge been analyzed following transplantation of stringently purified HSC^LT^, when all myeloid cells and their progenitors are truly HSC-derived. Surprisingly, GMP from pristane-exposed donor HSC^LT^ exhibited an overall reduction in chromatin accessibility relative to GMP from control HSC^LT^ across two independent biological replicates with 1213 unique peaks found across 1146 genes (**Fig 6B, Table S4**). When we examined the location of the differentially accessible regions (DAR), we found that all DAR were at sites of reduced chromatin accessibility and almost all were in gene promoters (92.0%) (**Fig 6C**). We then ran ORA on the DAR and strikingly, we found reduced chromatin accessibility at the promoters of genes associated with cellular metabolism, cell cycle, Myc and mTOR pathways (**Fig 6D**). Indeed, the *Mtor* locus itself exhibited reduced promoter accessibility in GMPs from pristane-exposed donor HSC^LT^ (**Fig 6E**). We found similar reductions in chromatin accessibility with other key glycolysis pathway genes such as *Gapdh* and *Hk2* (hexokinase 2). Using HOMER motif analysis, we found that the promoter sequences that were being altered were significantly enriched for binding sites for Sp2, Nfy, and Elf4 among others (**Fig S6A**). Functionally, Sp2 serves to recruit Nfy to DNA to transcribe genes necessary for metabolic activation (Ly, Yoshida and Yamaguchi, 2013; Völkel *et al*., 2015; Benatti *et al*., 2016), while Elf4 has been shown to limit the inflammatory response of macrophages (Tyler *et al*., 2021; Sun *et al*., 2022). Overall, these data suggest that HSC^LT^ from pristane-treated mice exhibit reduced chromatin accessibility at genes related to many metabolic processes.

**Figure 6.**
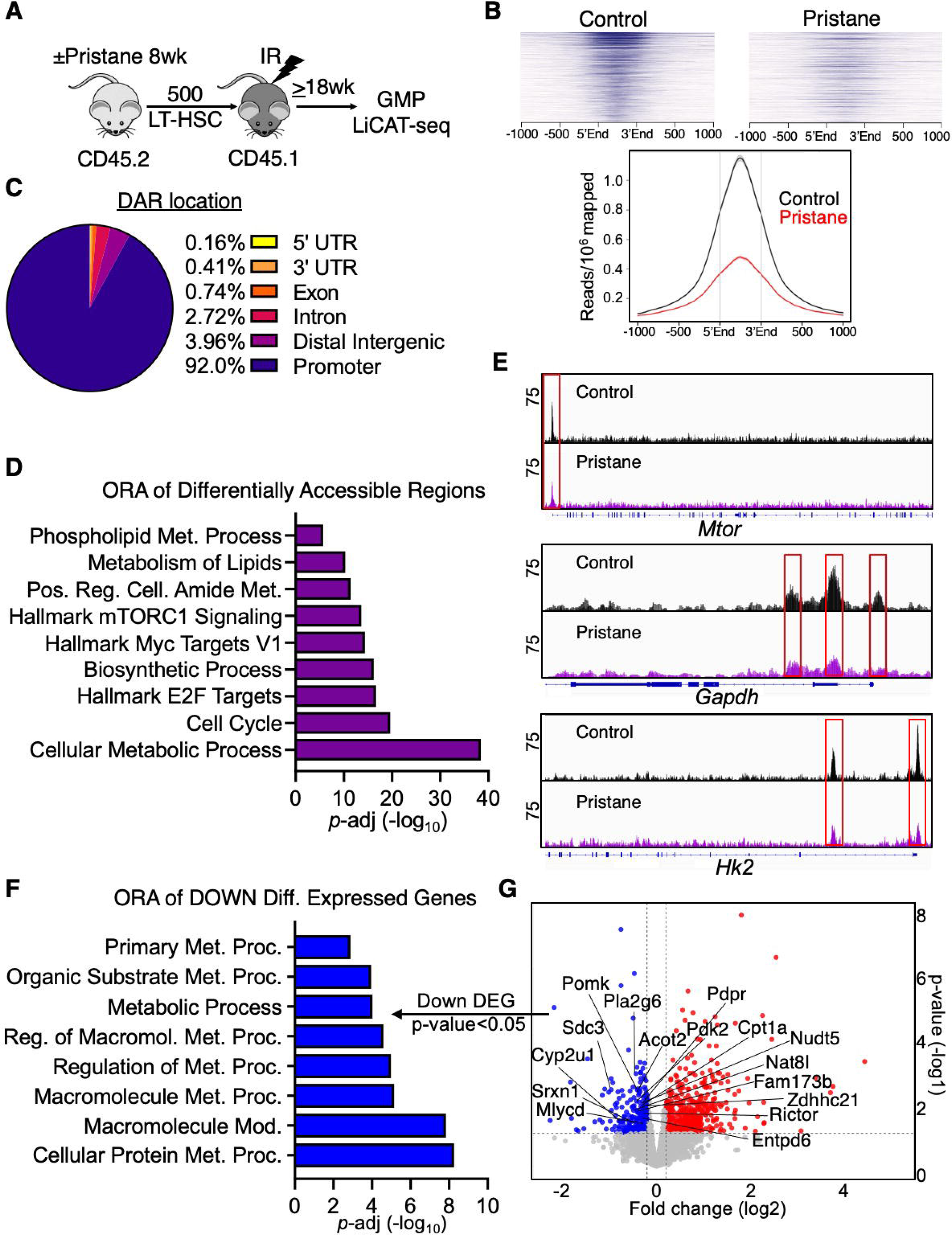
Pristane exposure induces heritable reprogramming of HSC^LT-^derived GMP. **(A-G)** Experimental design; mice were treated ± pristane for eight weeks, then 500 HSC^LT^ (LEK/Sca1^+^/Flk2^-^/CD48^-^/CD150^+^/EPCR^+^/CD34^-^) were FACS sorted from CD45.2 donor mice and then transplanted into lethally irradiated CD45.1 recipient mice. After 18 weeks of engraftment CD45.2 donor GMP were sorted for LiCAT-seq analysis; n = 2/group **(B)** Peak intensity heatmaps (top) from ATAC-seq analysis of CD45.2 donor derived GMP (From A), and quantification of reads mapped (bottom) **(C)** Location percentage of the differentially accessible regions (DAR) along the genome from ATAC-seq analysis of CD45.2 donor derived GMP (From A). **(D)** Over-representation analysis (ORA) of significantly enriched pathways (p-adj < 0.05) determined using the genes with significantly changed DAR (p-adj < 0.05) from the ATAC-seq analysis (From B). **(E)** Integrative Genomics Viewer (IGV) tracks of ATAC-seq read peaks of Mtor (top), Gapdh (middle), and HK2 (bottom); each track is an overlay of 2 biological replicates/group. **(F)** ORA of significantly enriched pathways (p-adj < 0.05) determined using the significantly down-regulated (p-val < 0.05) differentially expressed genes (DEG) from RNA-seq analysis of CD45.2 donor derived GMP (From A). **(G)** Volcano plot of significantly (p-value < 0.05) down-regulated (blue) and up-regulated (red) DEG from the RNA-seq analysis of CD45.2 donor derived GMP (From A). Key down-regulated genes are indicated.

We then evaluated transcriptional alterations in GMP derived from pristane-exposed donor HSC^LT^ by performing transcriptomic analysis of corresponding RNA pools generated by LiCAT-seq. To corroborate our ATAC-seq analysis, we first investigated which genes were downregulated in GMP. Unsurprisingly, numerous metabolic genes were significantly downregulated (p-val < 0.05) compared to control derived GMPs such as: *Cyp2u1* (hydroxylase metabolizing long chain fatty acids), *Acot2* (acyl-CoA hydrolase hydrolyzing acyl-CoA into fatty acids), *Cpt1a* (mitochondrial carnitine palmitase transporter), and notably *Rictor* (rapamycin resistant activator of mTOR) (**Fig 6G, Table S4**). Interestingly, many of the genes with reduced transcription, also showed reduced chromatin accessibility (**Fig S6B**). ORA analysis of the downregulated genes (pval<0.05) confirmed broad downregulation of metabolism pathways (**Fig 6F**). These data show that GMPs inherit an epigenetic program limiting chromatin accessibility to metabolic genes from pristane treated HSC^LT^, resulting in reduced transcription of metabolic pathway genes.

Since BMDMs derived from pristane-exposed donor HSC^LT^ exhibited features of immune training, we next investigated whether similar patterns of host defense and inflammatory genes identified in progenitor cells from other immune training models were similarly upregulated in GMP. Interestingly, GMP from pristane-exposed donor HSC^LT^ exhibited increased expression of *Jun*, *Junb*, *Jund*, *Fos, Fosb,* and *Atf3* transcription factors, which are all essential to activating and propagating proinflammatory gene programs (**Fig S6C**) (Schonthaler, Guinea-Viniegra and Wagner, 2011; Wang *et al*., 2012; Aung *et al*., 2013; Hannemann *et al*., 2017; Ding *et al*., 2020; Kim *et al*., 2021). To further evaluate the molecular priming of the GMP we ran ORA on upregulated DEG (p-val < 0.05; **Fig S6D**). Consistent with previous characterizations of immune training, we found that there was a significant enrichment in programs related to immune system activation, host defense responses and the AP-1 transcriptional complex, of which Jun and Fos are members. Importantly, recent work in epithelial stem cells have shown that Jun and Fos maintain trained immunity memory (Larsen *et al*., 2021). Overall, we find that prior pristane exposure imparts a unique epigenetic program in GMPs associated with suppression of cellular metabolism, alongside an activated transcriptional program priming cells for immune activation.

### BMDMs from pristane-exposed donor HSC^LT^ exhibit reduced glycolytic metabolism

We next investigated whether the reduction in metabolism predicted by LiCAT-seq was reflected functionally in HSC^LT^-derived BMDMs following exposure to chronic autoimmune inflammation. We again generated BMDMs from mice transplanted with pristane- or pIC-exposed donor HSC^LT^ (**Fig 7A**), stimulated them overnight with ImmCom and analyzed their metabolic properties using the Seahorse glycolytic and mitochondrial stress tests. (**Fig 7B, S7B**). Strikingly, BMDMs derived from pristane-exposed donor HSC^LT^ failed to increase glycolytic activity in response to ImmCom stimulation relative to control or pIC settings (**Fig 7C**). Likewise, only BMDMs derived from control donor HSC^LT^ were capable of triggering increased OXPHOS following ImmCom stimulation (**Fig S7C**). Overall, these data show that BMDMs derived from pristane-exposed HSC^LT^ exhibit reduced metabolic activity in response to immune stimulation and provide functional evidence in support of our LiCAT-seq data.

**Figure 7.**
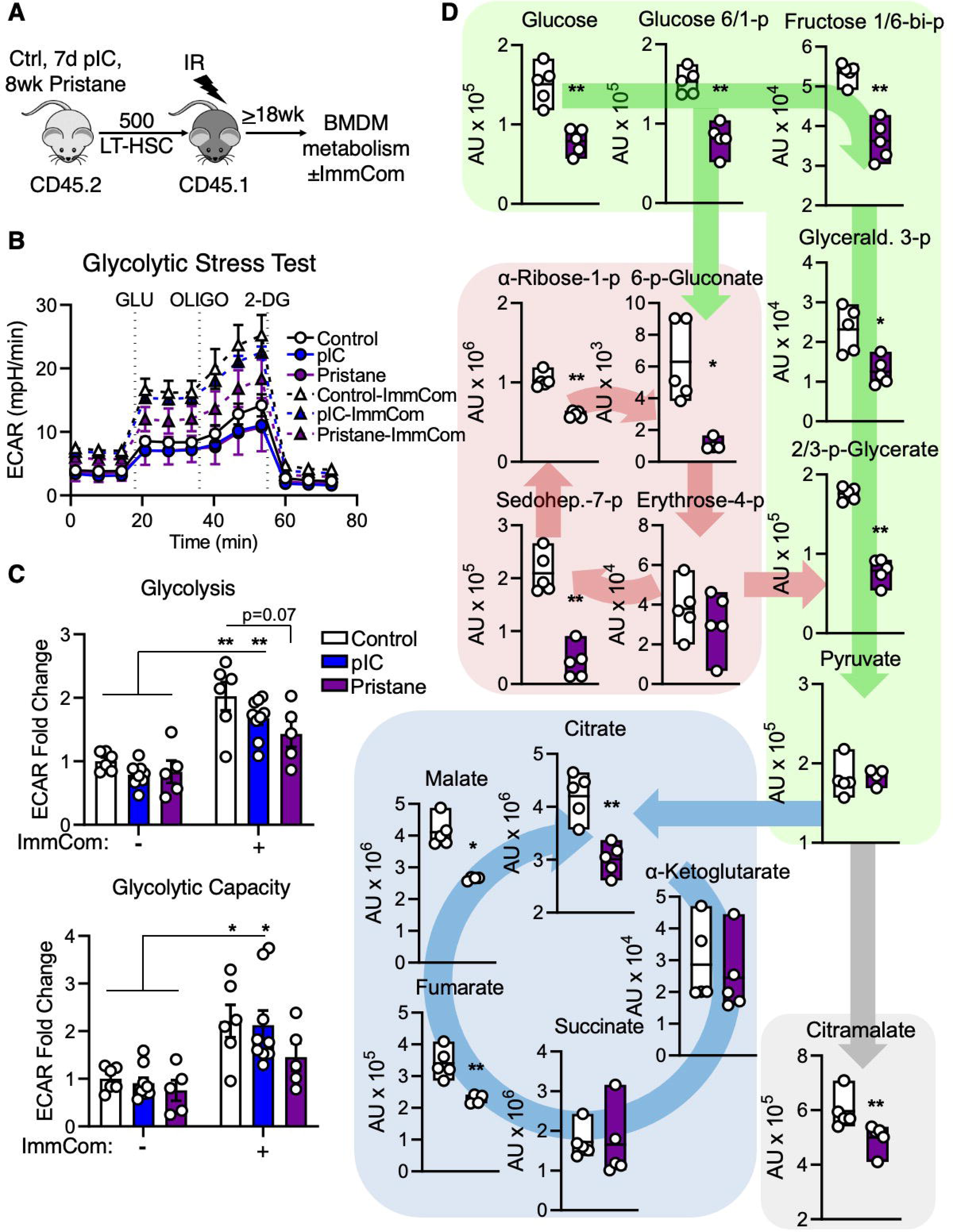
BMDMs from pristane-exposed donor HSC^LT^ exhibit reduced glycolytic metabolism. **(A-C)** Experimental design; mice were treated ± pristane for eight weeks or pIC for seven days, then 500 HSC^LT^ (LEK/Sca1^+^/Flk2^-^/CD48^-^/CD150^+^/EPCR^+^/CD34^-^) were FACS sorted from CD45.2 donor mice and then transplanted into lethally irradiated CD45.1 recipient mice. After 18 weeks of engraftment, bone marrow derived macrophages (BMDMs) were generated and then stimulated overnight with immune complexes (ImmCom) of IgG targeting NP-BSA_16_ before having their glycolytic metabolism assessed using the Seahorse glycolytic stress test. **(B)** Tracer of glycolytic stress test of unstimulated (solid lines) and ImmCom stimulated (dashed lines) BMDMs from mice transplanted with control, pIC-, or pristane-exposed donor HSC^LT^ (From A). **(C)** Quantification of glycolysis (ECAR after addition of the glucose (GLU); top), and glycolytic capacity (ECAR after the addition of oligomycin (OLIGO), bottom). Data are represented as fold change compared to Control unstimulated average ECAR; n = 6 Control, 9 pIC, 5 Pristane. **(D)** Mass spectrometry analysis of metabolites, after a six-hour stimulation with ImmCom, in BMDMs from mice transplanted with pristane-exposed or control donor HSC^LT^ (From A); n = 5/group. Data are represented as SEM. *p < 0.05, **p < 0.01, significance determined by Two-way ANOVA with Tukey’s post-test (C), or two-tailed Mann Whitney U-test (D).

To further characterize this metabolic state, we stimulated BMDMs for six hours with ImmCom prior to harvesting them for global metabolomics analysis using ultra-high performance liquid chromatography/mass spectrometry (UHPLC-MS) analysis. In agreement with our Seahorse data, BMDMs had an overall reduction in metabolites following ImmCom stimulation compared to control donor HSC^LT^-derived BMDMs (**Fig 7D**). This phenotype was particularly evident in several pathways constituting central carbon metabolism, including glycolysis (green), the pentose phosphate pathway (red), the tricarboxylic acid cycle (TCA, blue), as well as alternative carbon metabolites (grey). Strikingly, and consistent with a broad suppression of metabolic activity we identified global reductions in amino acids, nucleotides, glutathione intermediates, serine metabolism intermediates, and fatty acids among others (**Table S5**). Interestingly, there were also significant reductions in metabolites from central carbon metabolism, nucleotide metabolism, and fatty acids, etc even without ImmCom stimulation (**Fig S7D, Table S5**). Overall, these data show that BMDMs derived from pristane-exposed donor HSC^LT^ have reduced metabolic activity in response to ImmCom stimulation

## DISCUSSION

Herein, we demonstrate using rigorous transplantation-based approaches that HSC^LT^ constitute a long-term reservoir for trained myeloid cells in the setting of chronic autoimmune disease (AD). We show that BMDMs derived from these HSC^LT^ exhibit hallmarks of trained immunity, including increased cytokine secretion, bacterial killing activity, and upregulation of host defense gene programs in GMPs including expression of several AP-1 transcription factors previously identified as drivers of immune training (Larsen *et al*., 2021). Notably, we demonstrate that BMDMs derived from pristane-exposed donor HSC^LT^ are also capable of robustly activating T cells following stimulation with immune complexes (ImmCom). Lastly, we find that in contrast to BMDMs from mice with active AD, BMDMs derived from pristane-exposed donor HSC^LT^ exhibit attenuated metabolic activity following ImmCom stimulation, which is associated with reduced chromatin accessibility at the promoters of genes in key metabolic pathways associated with immune training, including glycolysis and mTOR (Cheng *et al*., 2014). Altogether, our data demonstrate that chronic inflammation associated with AD imprints a heritable program in HSC^LT^ that enhances the pro-inflammatory activity of downstream myeloid cells and potentiates their capacity to activate adaptive immune cells.

The observation that myeloid cells can retain enhanced inflammatory gene expression patterns long past their individual lifetimes has implicated longer-lived hematopoietic stem and progenitor cells (HSPC) as the most likely reservoirs for immune training (Garly *et al*., 2003). To address this point, previous studies have used transplantation of unfractionated BM cells or heterogenous HSPC-enriched BM fractions, such as the phenotypic LSK compartment, to evaluate the contribution of HSPC to immune training. (Kaufmann *et al*., 2018; de Laval *et al*., 2020; Kalafati *et al*., 2020). Here, to rigorously evaluate whether HSC^LT^ themselves are a sufficient functional reservoir for immune training, we used transplantation of purified EPCR^+^/CD34^-^ SLAM cells, which we previously showed to be highly enriched for HSC^LT^ activity under chronic inflammatory stress conditions (Rabe *et al*., 2020). Indeed, we find that BMDMs and GMPs derived from these HSC^LT^ exhibit well-characterized hallmarks of immune training, including increased expression of transcriptional regulators Jun and Fos in GMPs. These data support a role for HSC^LT^ themselves as key players in immune training. In contrast to previous work using canonical approaches to induce immune training *in vivo* such as single injections of BCG, β-glucan and/or LPS, we sought to address whether exposure to chronic AD-associated inflammation could likewise impart a heritable ‘trained’ phenotype in HSC^LT^ and their myeloid progeny. Interestingly, BMDMs from pristane-exposed HSC^LT^ exhibit more robust evidence of immune training relative to BMDMs from HSC^LT^ exposed to 7d of pIC, which we used as a short-term inflammatory stimulation that does not trigger an autoimmune cascade. Both stimuli have been shown to activate broadly similar systemic cytokine profiles, most notably IFN-Is. Interestingly, IFN signaling programs are commonly found in RNA-seq analysis of SLAM cells and HSPC from mice stimulated with BCG or -glucan (Netea, Joosten and van der Meer, 2017; Kaufmann *et al*., 2018; Chavakis, Mitroulis and Hajishengallis, 2019; Cirovic *et al*., 2020; Kalafati *et al*., 2020), further supporting the role of IFNs in mediating trained immunity. However, we suspect length of exposure may be a contributing factor to the more robust induction of immune training in BMDMs from pristane-exposed donor HSC^LT^. However, is not yet well understood how inflammatory activity imparts epigenetic memory on deeply quiescent HSC^LT^ in an *in vivo* setting, either through direct signaling to HSC^LT^ themselves or via indirect mechanisms such as alteration of the BM microenvironment. Furthermore, it is not understood whether immune training is imparted on all HSC following an inflammatory stimulus or only a subset of clone(s) defined by dormancy level, location in the BM niche, cytokine receptor expression, and/or other factors. With this in mind, the more robust trained phenotype of BMDMs derived from pristane-exposed HSC^LT^ could be the result of either more HSC clones undergoing epigenetic remodeling over time, or a reinforcement of epigenetic remodeling in certain susceptible HSC clone(s). Clonal analyses will be needed to better understand the dynamics of this process, including whether HSC ‘training’ is a clonal process, and what factor(s) such as time, localization in the BM, underlying gene expression and metabolic activity, and/or cell cycle activity dictate whether an HSC clone is susceptible to epigenetic remodeling to retain ‘memory’ of an inflammatory insult.

Interestingly, we find that BMDMs derived from pristane- and pIC-exposed HSC^LT^ exhibit a novel increase in their capacity to present antigen to T cells following stimulation with ImmCom. In the setting of AD, ImmCom are crucial drivers of disease pathology, and their deposition in the kidney leads to glomerulonephritis, a severe morbidity in SLE patients (Satoh *et al*., 1995; Herrada *et al*., 2019). Further, expression of FcγR on myeloid cells from SLE patients positively correlates with disease severity, and incidence of lupus nephritis (Li *et al*., 2009; Li *et al*., 2010; Abd-Elhamid *et al*., 2017), highlighting the unique relationship between ImmCom and myeloid cells in AD pathogenesis. Importantly, myeloid cells are becoming increasingly appreciated for their role in triggering the autoimmune cascades that initiate AD (Laria *et al*., 2016; Morell, Varela and Marañón, 2017; Navegantes *et al*., 2017; Ushio *et al*., 2017; Ma *et al*., 2019). Indeed, the augmented T cell co-stimulatory activity combined with increased production of inflammatory cytokines we observe following ImmCom stimulation of BMDMs derived from pristane-exposed HSC^LT^ could represent a maladaptive phenotype capable of initiating bystander activation of autoimmune T cell clones (Nogai *et al*., 2005; Fujinami *et al*., 2006; Pacheco *et al*., 2019). It has recently been shown that inflammation from ligature induced periodontitis can durably train SLAM-HSC, and post transplantation generate hyperinflammatory myeloid cells which increase the severity of collagen antibody-induced arthritis in recipient mice (Li *et al*., 2022), highlighting the relationship between maladaptive TI and AD development. This maladaptive process may also underlie the remitting-relapsing course of AD like SLE, wherein HSC^LT^ serve as a pathogenic reservoir for trained myeloid cells that are capable of reactivating autoimmune cells in response to a physiological insult, subsequently triggering characteristic disease ‘flares’ in individuals whose disease is otherwise in remission. Along these lines, previous work analyzing the transcriptomes of PBMCs from SLE patients found that monocytes retained a hyper-inflammatory gene expression phenotype even in patients whose disease was in remission (Labonte *et al*., 2018). Notably, the transcriptomes of B and T cells from the same patients returned to normal during remission, supporting a model in which long-term heritable changes associated with immune training in HSC may be uniquely expressed in their short-lived myeloid progeny. Furthermore, previous investigations have established a correlative relationship between AD and prior infection, suggesting immune training resulting from the earlier insult acts as an etiological driver of AD. Further investigations can establish which, if any, epigenetic alterations are passed on to lymphoid cells from HSC, and if these changes have functional consequences for AD development. In addition, the extent to which prior inflammatory insults modulate AD pathogenesis, and the participation of HSC in this process, remain open questions to be addressed.

Activation of mTOR and glycolysis are critical metabolic pathways needed to support the establishment of TIin myeloid cells (Cheng *et al*., 2014; Arts *et al*., 2016) In line with these studies, we find that monocytes and GMPs isolated directly from primary pristane-treated mice exhibit upregulation of glycolysis and OXPHOS metabolic pathways along with induction of gene programs associated with cellular activation. BMDMs from these mice (and to a lesser extent, pIC-treated mice) exhibit classic hallmarks of trained immunity, including a significant upregulation of glycolytic metabolism following stimulation with ImmCom. Thus, we were surprised to find that BMDMs derived from pristane-exposed donor HSC^LT^ exhibited attenuated metabolic activity following ImmCom stimulation, as measured by extracellular flux and mass spectrometry-based measurement of glycolysis and TCA metabolites. Strikingly, this phenotype was accompanied by a specific reduction in chromatin accessibility and expression of genes associated with glycolysis, mTOR and cell cycle activity in GMP derived from pristane-exposed donor HSC^LT^. These data suggest that glycolytic activity is clearly coupled to the establishment of immune training, as evaluated in short-term assays using BMDMs or PBMCs (Bekkering *et al*., 2016, 2018; Arts *et al*., 2016). On the other hand, once the epigenetic program for immune training is established, glycolysis may become dispensable either as a source of energy for induction of inflammatory gene programs and/or as a source for catabolic derivatives used for epigenetic modifications, like acetyl-CoA and/or TCA cycle intermediates that tune the activity of epigenetic writers and/or erasers (Fawal and Davy, 2018; Domínguez-Andrés *et al*., 2019; Yang and Jiang, 2020). Hence, trained myeloid cells derived from these HSC may either come to rely on other sources of energy (amino/fatty acids, autophagy) for enhanced inflammatory and immune function. Flux assays to track the uptake and fate of glucose could offer further insight into how it is used differently by BMDMs derived from primary BM vs. transplanted HSC^LT^. Lastly, it is possible that the unique metabolic phenotype of BMDMs derived from pristane-exposed donor HSC^LT^ may reflect stress response mechanisms triggered by chronic exposure to inflammatory stress, which we and others have shown serves to limit the metabolic activity of HSC^LT^. Hence, glycolysis- and mTOR-dependent induction of immune training in the HSC compartment could be linked to the transient cell cycle induction triggered by acute inflammation, when key metabolic drivers like Myc, mTOR and glycolytic activity are briefly activated (Ito and Suda, 2014; Meng, Frank and Jewell, 2018). Conversely, long-term inflammatory exposure and/or changes in the systemic availability of metabolites could induce heritable activation of stress pathways via TFEB or other regulators that restrict metabolism to enforce dormancy (García-Prat *et al*., 2021). Our transplantation assays could potentially be seen as an accelerator of clonal succession wherein we force HSC to undergo replication, thereby selecting for long-lived HSC clone(s) with reduced mTOR/glycolysis activity following induction of trained immunity. Further studies can address the extent to which increased metabolic activity via glycolysis and mTOR is required to establish epigenetic patterns associated with immune training in HSC^LT^.

Altogether, our data demonstrate that chronic inflammation associated with AD serves as a driver of immune training, with HSC^LT^ serving as the reservoir. This mechanism could establish a vicious cycle of chronic inflammation and maladaptive TIat the heart of AD development and relapse.

## Acknowledgments

We thank Garrett Hedlund for expert assistance with flow cytometry resources. We also thank Rachel Gessner, James Chavez and Katia Niño for additional experimental support. This work was supported by K01 DK098315, the Cleo Meador and George Ryland Scott Endowed Chair in Hematology, and CU Department of Medicine Outstanding Early Career Scholars Program (to E.M.P.), R35 HL155672 (to K.Y.K.), the National Science Foundation Graduate Research Fellowship Program (to T.S.M.), R01 DK125595 (to M.A.B.), R01 HL146442 (to A.D), and T32 DK060445 (to B.K.). We would also like to thank Raul Torres for his generous gift of purified anti-NP-BSA_16_ used for immune complex generation.

## Author Contributions

Conceptualization: T.S.M. and E.M.P; Methodology: T.S.M., B.K., M.A.B., E.D., R.C.H, A. G., J.R.R., A.D., K.Y.K., and E.M.P.; Investigation: T.S.M., B.K., M.A.B., E.D., R.C.H., A. G., A.D., K.Y.K., and E.M.P.; Resources: J.R.R.; Writing – Original Draft: T.S.M. and E.M.P.; Writing – Review and Editing: T.S.M., B.K., M.A.B., K.Y.K., E.M.P.; Supervision: K.Y.K and E.M.P.; Funding Acquisition: K.Y.K. and E.M.P.

## Supplemental Figure Legends

**Figure S1, related to figure 1.**
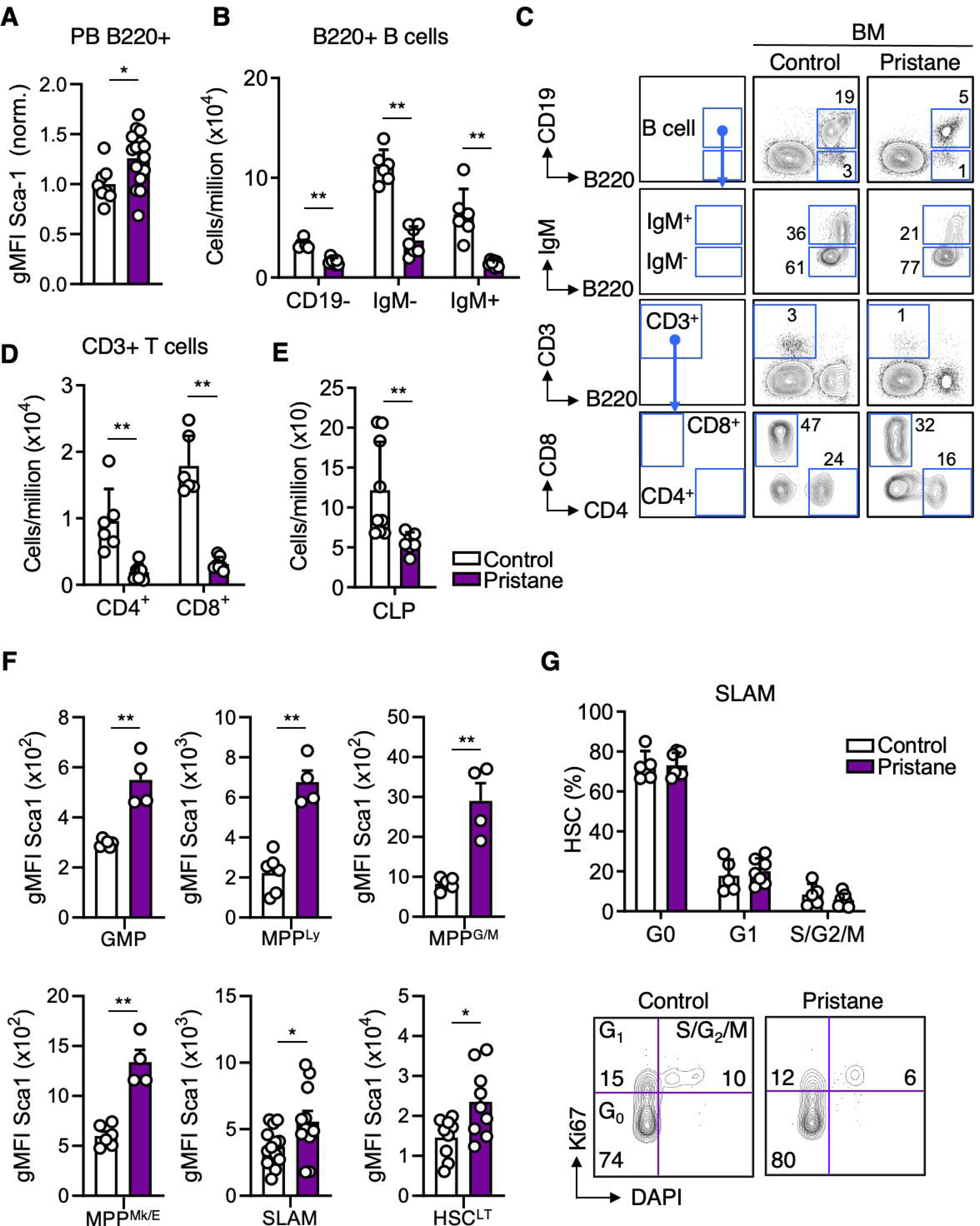
Experimental design: mice were treated ± pristane for eight weeks for FACS analysis. **(A)** Flow cytometry analysis of geometric mean fluorescent intensity (gMFI) of Sca-1 on PB B220+ lymphocytes; n = 8 Control, 16 Pristane. **(B-D)** Quantification of FACS analysis of (B) B220+ B cell populations and (C) representative FACS plots for mature bone marrow lymphocyte populations, and (D) quantification of CD3+ T cell populations; n = 6/group **(E)** Quantification of FACS analysis of common lymphoid progenitors (CLP) in the bone marrow; n = 9 Control, 7 Pristane. **(F)** Flow cytometry analysis of geometric mean fluorescent intensity (gMFI) of Sca-1 on BM hematopoietic progenitor populations; n = 6-13 Control, 4-11 Pristane. **(G)** Quantification of the frequency (Top) and representative FACS plots (bottom) of cell cycle distribution of SLAM cells; n = 5 Control, 7 Pristane Data are represented as SEM. *p < 0.05, **p < 0.01, significant versus control; significance determined by two-tailed Mann Whitney U-test.

**Figure S2, related to figure 1.**
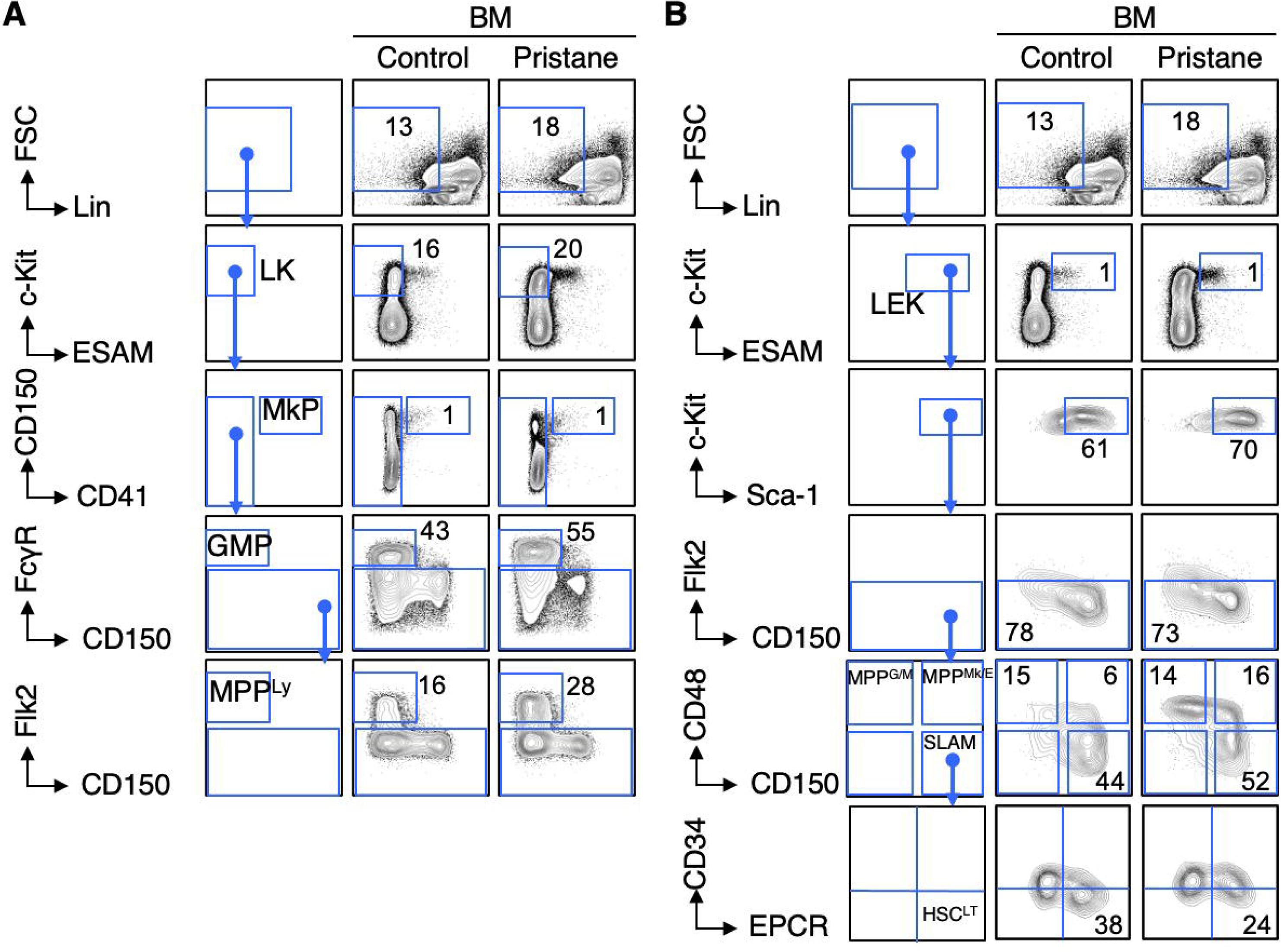
**(A-B)** Representative FACS plots and gating strategy for (A) myeloid progenitors and MPP^LY^, and (B) immature hematopoietic stem and progenitor cells from the bone marrow of mice treated ± pristane for eight weeks

**Figure S3, related to figure 2 and 3.**
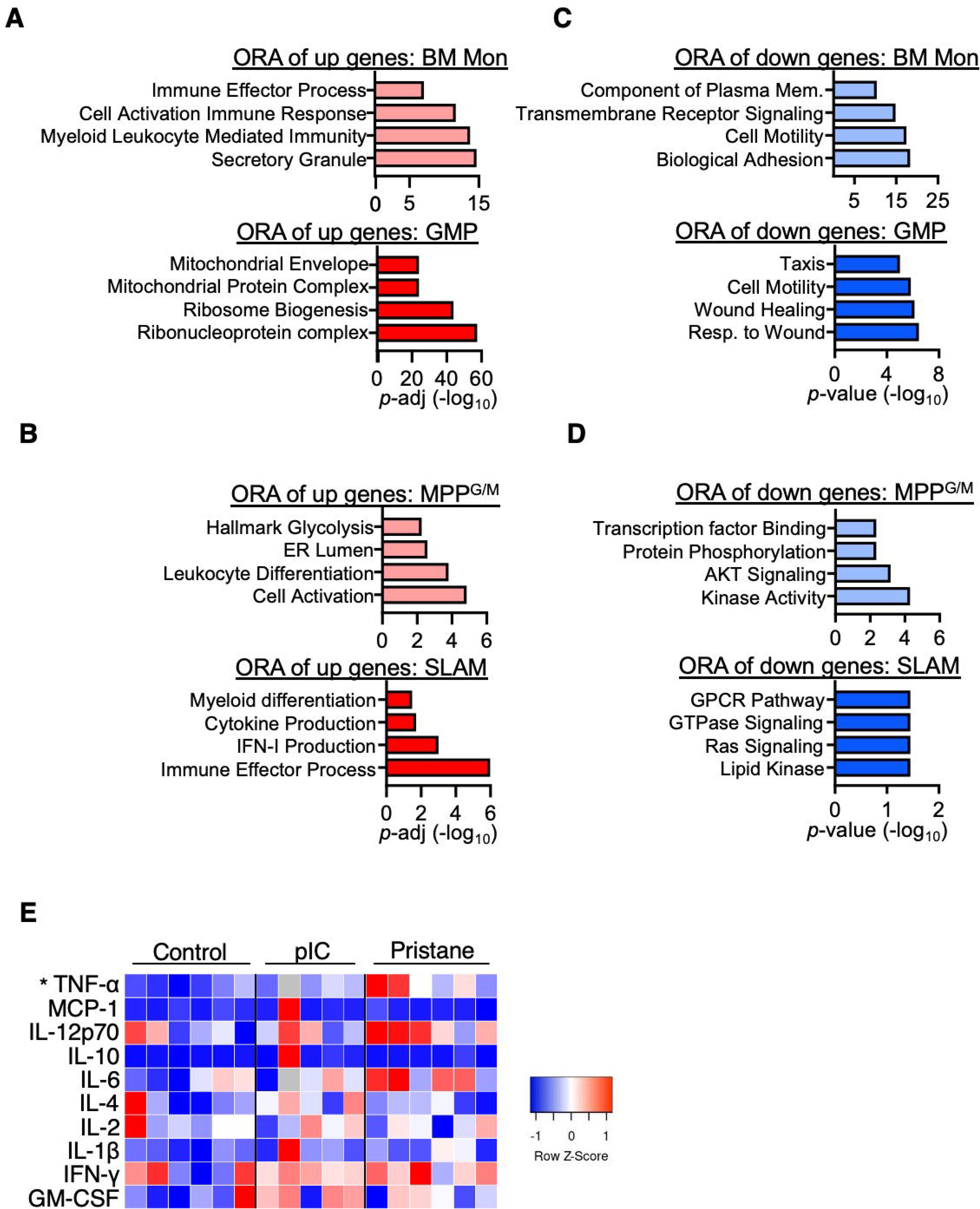
Experimental design: mice were treated ± pristane for four weeks for RNA-seq analysis; n = 3/group **(A-B)** Over-representation analysis (ORA) of significantly enriched pathways (p-adj < 0.05) determined using the significantly up-regulated (p-adj < 0.05) differentially expressed genes (DEG) (From Fig 2C) in (A) BM Mon (top) and GMP (bottom), and (B) MPP^G/M^ (top) and SLAM (bottom). **(C-D)** Over-representation analysis (ORA) of significantly enriched pathways (p-adj < 0.05) determined using the significantly down-regulated (p-adj < 0.05) DEG (From Fig 2C) in (C) BM Mon (top) and GMP (bottom), and (D) MPP^G/M^ (top) and SLAM (bottom). **(E)** (Related to Fig 3D) Heatmap of cytokines in cell culture supernatant at the end of the 72hr killing assay (From 3B); significance to control; n = 6 Control, 5 pIC, 6 Pristane. Significance determined by One-way ANOVA with Tukey’s post-test. =

**Figure S4, related to figure 4.**
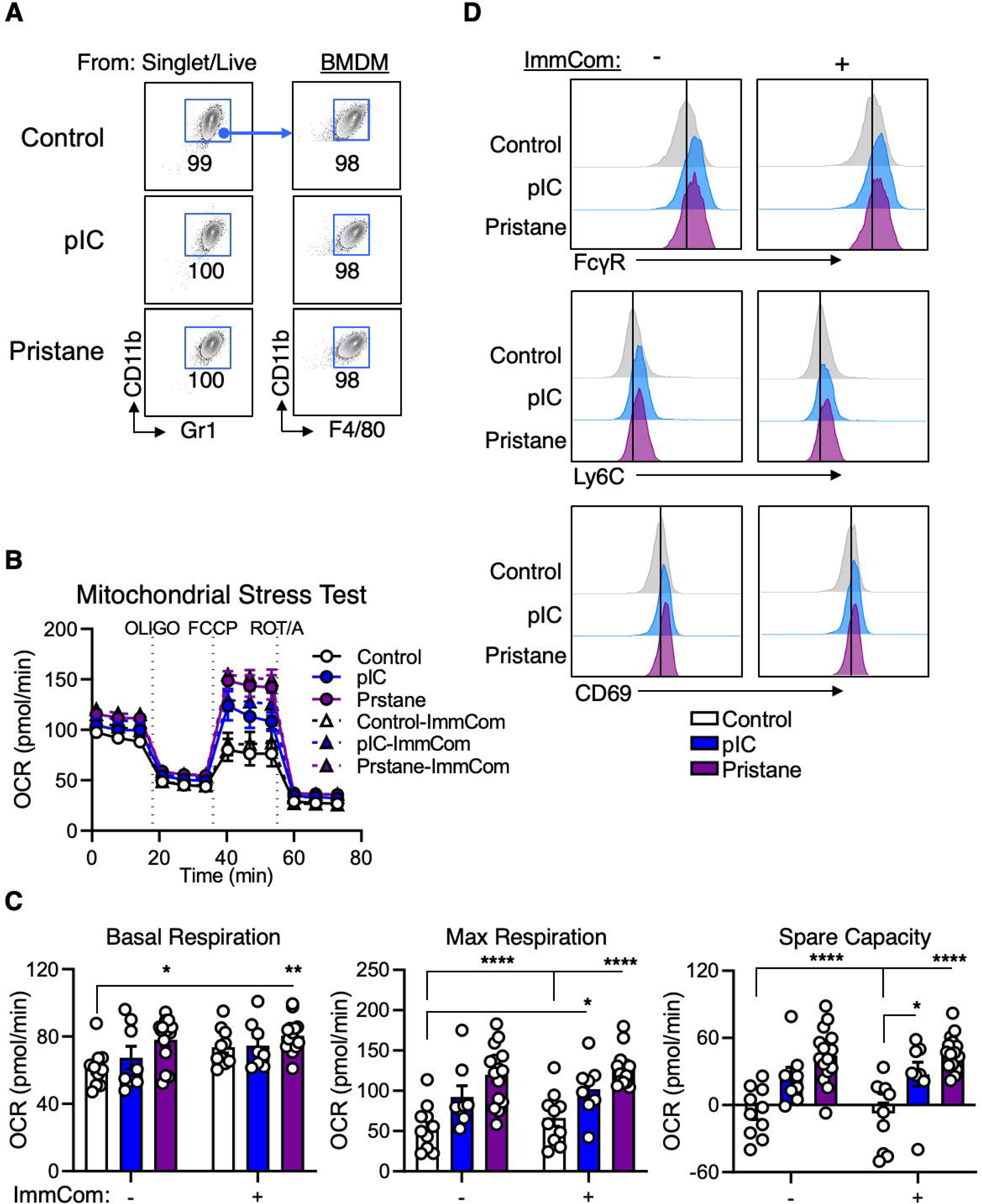
Experimental design: BMDMs were generated from mice that were treated ± pristane for eight weeks or pIC for seven days, and then stimulated with immune complexes (ImmCom) of IgG targeting NP-BSA_16_. **(A)** Representative FACS plots and gating strategy for identification of bone marrow derived macrophages (BMDMs). **(B)** Tracer of mitochondrial stress test of BMDMs that were unstimulated (solid lines) or stimulated overnight with ImmCom (dashed lines); n = 14 Control, 8 pIC, 15 Pristane. **(C)** Quantification of basal respiration (OCR after the addition of oligomycin (Oligo); left), maximal respiration (OCR after the addition of FCCP, middle), and spare capacity (the difference in OCR between basal and maximal respiration, right) in BMDMs that were stimulated overnight with ImmCom (From B); n = 14 Control, 8 pIC, 15 Pristane. **(D)** Representative histograms of surface protein expression on BMDM from mice treated with pristane for eight weeks, pIC for seven days, or control that were then stimulated with ImmCom for 24hr (FcγR left, and Ly6C middle) or 72hr (CD69 right); n = 10 Control, 9 pIC, 11 Pristane. Data are represented as SEM. *p < 0.05, **p < 0.01, ***p < 0.001, ****p < 0.0001; significance determined using Two-way ANOVA with Tukey’s post-test.

**Figure S5, related to figure 5.**
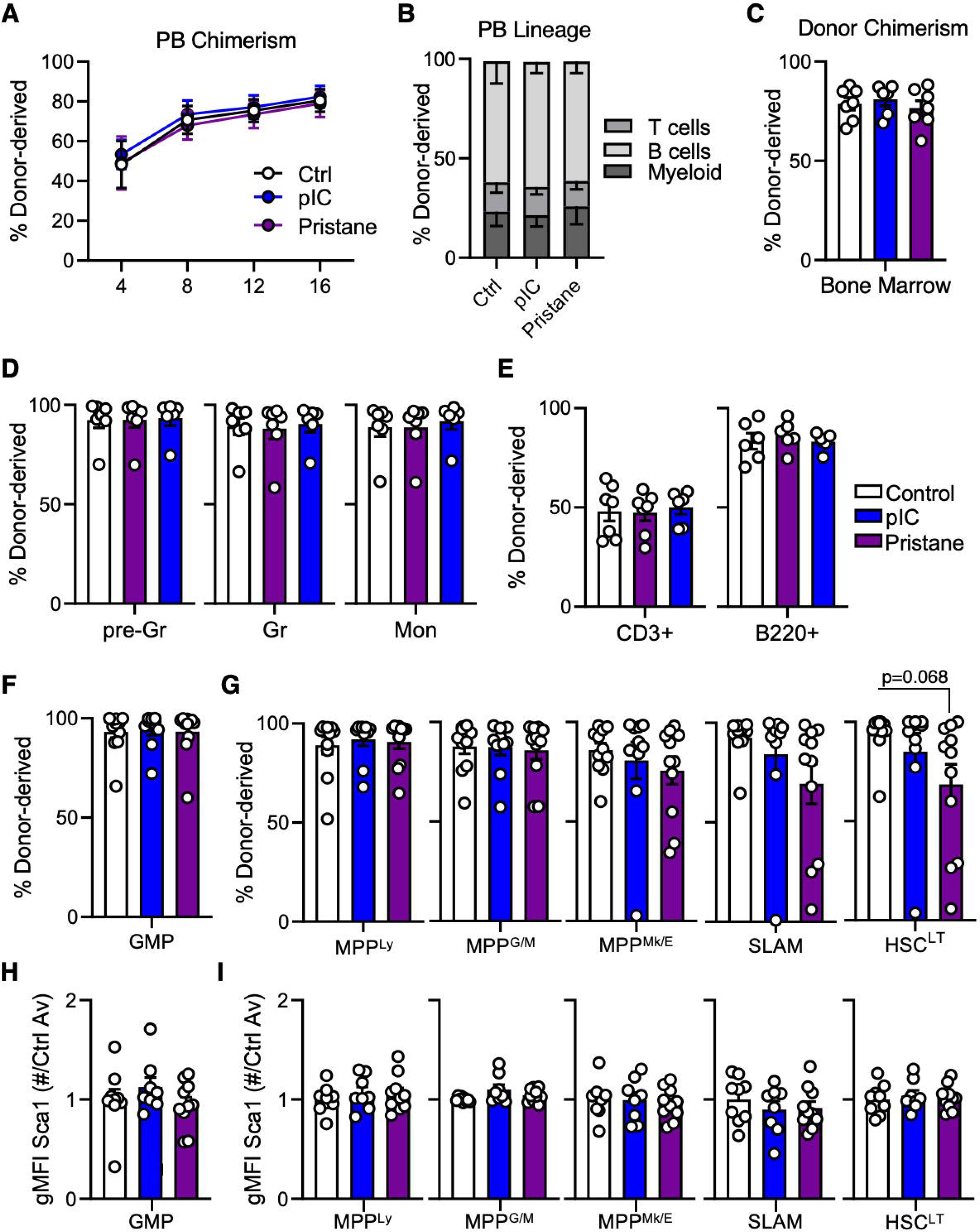
Experimental design: CD45.2 donor mice were treated ± pristane for eight weeks or pIC for seven days, then 500 HSC^LT^ (LEK/Sca1^+^/Flk2^-^/CD48^-^/CD150^+^/EPCR^+^/CD34^-^) were FACS sorted and transplanted into lethally irradiated CD45.1 recipient mice. **(A-B)** (A) Percent of donor chimerism over time and (B) the donor lineage distribution in the peripheral blood (PB) of mice transplanted with control, pIC-, or pristane-exposed donor HSC^LT^ after 16 weeks of engraftment; n = 17 Control, 16 pIC, 19 Pristane. **(C)** Percent of donor CD45.2 chimerism in the bone marrow (BM) of mice transplanted with HSC^LT^ from control, pIC-, or pristane-treated donor mice after 18 weeks of engraftment; n = 17 Control, 16 pIC, 19 Pristane. **(D-G)** Quantification of donor chimerism using FACS analysis in: (D) mature myeloid cells, (E) mature lymphocytes, (F) myeloid progenitors, and (G) immature hematopoietic stem and progenitor cells; n = 11 Control, 10 pIC, 11 Pristane. **(H-I)** Flow cytometry analysis of geometric mean fluorescence intensity (gMFI) of Sca-1 on (H) granulocyte monocyte progenitors (GMPs), and (I) immature hematopoietic stem and progenitor cells; n = 9 Control, 8 pIC, 12 Pristane.

**Figure S6, related to figure 6:**
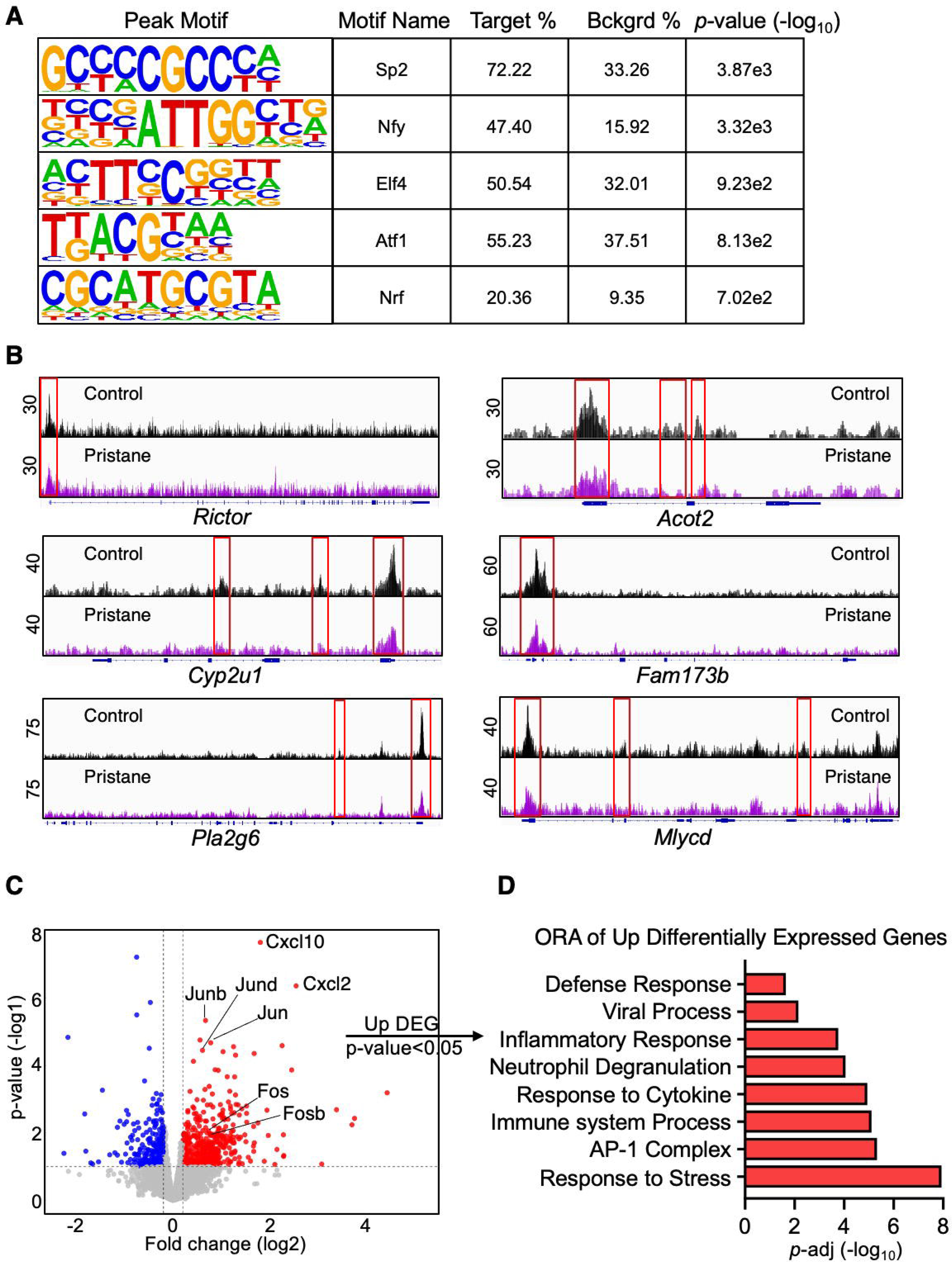
Experimental design: CD45.2 donor mice were treated ± pristane for eight weeks then 500 HSC^LT^ (LEK/Sca1^+^/Flk2^-^/CD48^-^/CD150^+^/EPCR^+^/CD34^-^) were FACS sorted and transplanted into lethally irradiated CD45.1 recipient mice. After 18 weeks of engraftment CD45.2 donor GMP were sorted for LiCAT-seq analysis; n = 2/group **(A)** Homer motif analysis of the significantly altered differentially accessible regions (DAR) showing the DNA motif associated with the peak, the associated transcription factor, the percent of the DAR with this motif, the amount of background signal, and the significance. **(B)** Integrative Genomics Viewer (IGV) tracks of ATAC-seq read peaks of metabolic genes identified in the RNA-seq analysis as significantly down-regulated (p-val < 0.05) in CD45.2 donor GMP (From Fig 6G); each track is an overlay of 2 biological replicates/group. **(C)** Volcano plot of significantly (p-value<0.05) down-regulated (blue) and up-regulated (red) differentially expressed genes (DEG) from the RNA-seq analysis of CD45.2 donor derived GMP. Key up-regulated genes are indicated. **(D)** Over-representation analysis (ORA) of significantly enriched pathways (p-adj < 0.05) determined using the significantly up-regulated (pval < 0.05) DEG from the RNA-seq analysis of CD45.2 donor derived GMP.

**Figure S7, related to figure 7:**
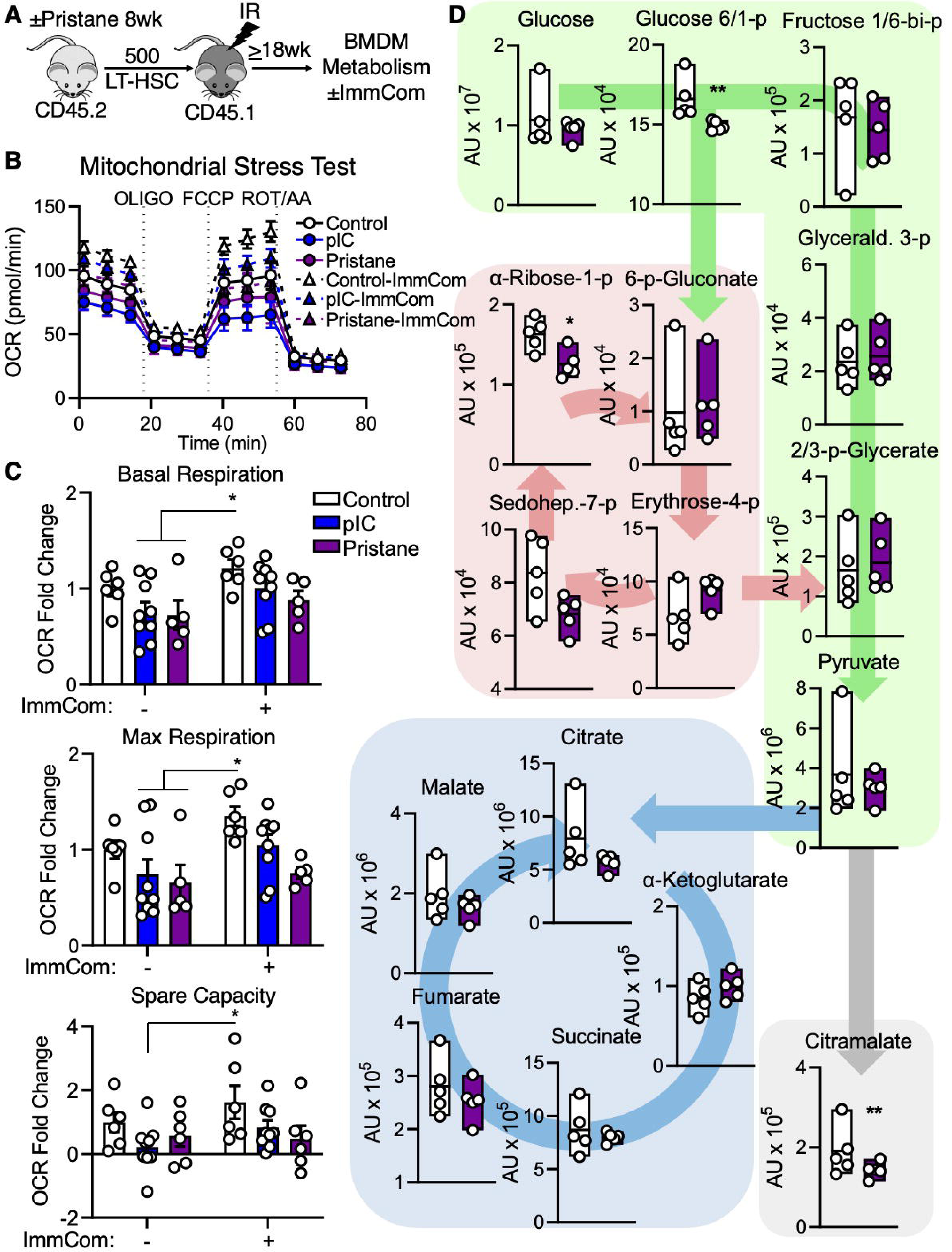
**(A-C)** Experimental design: mice were treated ± pristane for eight weeks or pIC for seven days, then 500 HSC^LT^ (LEK/Sca1^+^/Flk2^-^/CD48^-^/CD150^+^/EPCR^+^/CD34^-^) were FACS sorted from CD45.2 donor mice and transplanted into lethally irradiated CD45.1 recipient mice. After 18 weeks of engraftment bone marrow derived macrophages (BMDMs) were generated and then stimulated overnight with immune complexes (ImmCom) of IgG targeting NP-BSA_16_ before having their mitochondrial metabolism assessed using the Seahorse mitochondrial stress test. **(B)** Tracer of mitochondrial stress test of unstimulated (solid lines) and ImmCom stimulated (dashed lines) BMDMs from mice transplanted with control, pIC- or pristane-exposed donor HSC^LT^ (From A); n = 6 Control, 9 pIC, 5 Pristane. **(C)** Quantification of basal respiration (OCR after the addition of oligomycin (Oligo); top), maximal respiration (OCR after the addition of FCCP, middle), and spare capacity (the difference in OCR between basal and maximal respiration, bottom) in BMDMs that were stimulated overnight with ImmCom (from B). Data are represented as fold change compared to Control unstimulated average OCR; n = 6 Control, 9 pIC, 5 Pristane. **(D)** Mass spectrometry analysis of metabolites in unstimulated BMDMs from mice transplanted with control or pristane-exposed donor HSC^LT^ (From A); n = 5/group. Data are represented as SEM. *p < 0.05, **p < 0.01, significance determined by Two-way ANOVA with Tukey’s post-test (C), or two-tailed Mann Whitney U-test (D).

